# *BCM3D 2.0*: Accurate segmentation of single bacterial cells in dense biofilms using computationally generated intermediate image representations

**DOI:** 10.1101/2021.11.26.470109

**Authors:** Ji Zhang, Yibo Wang, Eric D. Donarski, Tanjin T. Toma, Madeline T. Miles, Scott T. Acton, Andreas Gahlmann

## Abstract

Accurate detection and segmentation of single cells in three-dimensional (3D) fluorescence timelapse images is essential for observing individual cell behaviors in large bacterial communities called biofilms. Recent progress in machine-learning-based image analysis is providing this capability with every increasing accuracy. Leveraging the capabilities of deep convolutional neural networks (CNNs), we recently developed bacterial cell morphometry in 3D (*BCM3D*), an integrated image analysis pipeline that combines deep learning with conventional image analysis to detect and segment single biofilm-dwelling cells in 3D fluorescence images. While the first release of *BCM3D* (*BCM3D 1*.*0*) achieved state-of-the-art 3D bacterial cell segmentation accuracies, low signal-to-background ratios (SBRs) and images of very dense biofilms remained challenging. Here, we present *BCM3D 2*.*0* to address this challenge. *BCM3D 2*.*0* is entirely complementary to the approach utilized in *BCM3D 1*.*0*. Instead of training CNNs to perform voxel classification, we trained CNNs to translate 3D fluorescence images into intermediate 3D image representations that are, when combined appropriately, more amenable to conventional mathematical image processing than a single experimental image. Using this approach, improved segmentation results are obtained even for very low SBRs and/or high cell density biofilm images. The improved cell segmentation accuracies in turn enable improved accuracies of tracking individual cells through 3D space and time. This capability opens the door to investigating time-dependent phenomena in bacterial biofilms at the cellular level.

## Introduction

Most terrestrial bacteria live in extended 3-dimensional tissue-like communities, named biofilms. As multicellular communities, bacteria can successfully colonize various biotic and abiotic surfaces. Biofilm-dwelling bacteria interact intimately not only with each other and the surface they reside on, but also with a self-produced extracellular matrix (ECM) that consists of proteins, DNA, and polysaccharides^1-3^. The sum total of these interactions helps biofilms develop emergent capabilities beyond those of isolated cells^1, 2, 4, 5^. Most notably, biofilms are more tolerant towards physical, chemical, and biological stressors^5-7^. Understanding how such capabilities emerge from the cooperative or antagonistic behaviors among individual cells requires live-cell compatible imaging technologies that are capable of resolving and tracking single cells within dense 3D biofilms.

Recently developed light sheet-based fluorescence imaging modalities combine high resolution with fast imaging speed and low phototoxicity at levels that cannot be matched by confocal microscopy^8-10^. Light sheet-based microscopy modalities are therefore increasingly used for non-invasive time-lapse imaging of eukaryotic cells and tissues^11-13^ as well as bacterial biofilms^14-16^. Depending on the type of biofilm, the cell density may however be too high to clearly resolve the gaps between cells with diffraction-limited microscopy. Super-resolution imaging modalities, such as structured illumination microscopy^17, 18^, improve the spatial resolution, but experimental improvements in spatial resolution come at the cost of decreased temporal resolution and increased light exposure to the specimen, which again raises photobleaching and phototoxicity concerns^19, 20^. An additional challenge arises for cell tracking studies. Tracking motile cells may require high frame rate imaging to achieve sufficient temporal resolution. Higher frame rates need to be accompanied by a proportional decrease in excitation laser intensities to mitigate photobleaching and phototoxicity. The decreased excitation laser intensities then result in lower signal-to-background ratios (SBRs) and signal-to-noise ratios (SNRs) in the individual images. The inherent trade-offs between spatial and temporal resolution, SBR/SNR, and photobleaching and phototoxicity is driving the continued development of new and improved image processing approaches that extract ever increasing amounts of useful information from the available experimental images.

Image processing pipelines based on supervised training of deep convolutional neural networks (CNNs) have been shown to outperform conventional image processing approaches for a variety of tasks in biomedical image analysis^21, 22^. For 3D biofilm image segmentation, we have recently developed Bacterial Cell Morphometry 3D (*BCM3D 1*.*0*), which achieved state-of-the-art performance for bacterial cell counting and cell shape estimation^23^. *BCM3D 1*.*0* does not rely on manually annotated training data, but instead combines *in silico*-trained CNNs for voxel classification with graph-theoretical linear clustering (mLCuts^24^) to post-process the thresholded CNNs outputs (i.e. the confidence maps for voxel-level classification). Using this approach, *BCM3D 1*.*0* automatically identifies individual cells in 3D images of 3D bacterial biofilms, reports their 3D shape and orientation, and classifies cell types with different morphologies. However, processing images with low SBRs and high cell densities remains challenging. Specifically, over- and under-segmentation errors increase in frequency for low SBR and high cell density images.

*Cellpose*^25^, *StarDist*^26^ and the work by Scherr *et al*.,^27^ are CNN-based approaches that create intermediate image representations for better segmentation. We reasoned that solving an image-to-image translation task may prove to be a more robust strategy for handling extreme imaging conditions than the voxel classification approach implemented in *BCM3D 1*.*0* or, at least, yield complementary segmentation results to *BCM3D 1*.*0*. Two different intermediate image representations are generally employed. The first representation is used to locate objects and the second representation is used to highlight the boundaries of objects. In previous work^25-27^, the CNN-predicted Euclidean distance to the nearest background pixel/voxel or the CNN-predicted object/background probability map was used to locate objects. Generation of boundary representations vary more widely: *StarDist* and *Cellpose* use star-convex polygons and spatial gradients separately to give complete boundaries, which can be used for object shape estimation. Scherr *et al*. instead enhance boundary regions that are close to other objects to prevent them from merging. Inspired by these approaches, we expanded the *BCM3D* workflow with a complementary CNN-based processing pipeline that translates the raw 3D fluorescence images into two distinct intermediate image representations that, in combination, are more amenable to conventional mathematical image processing, namely seeded watershed^28^ and Otsu thresholding^29^. For object localization, we adapted the approach used by *StarDist*^30^ and Scherr et al.^27^. For boundary information, however, we generated a new intermediate image representation that provides a complete 3D boundary of an object and additionally highlights whether the boundary is near other objects. We establish that, when combined and processed appropriately, these intermediate image representations provide biofilm segmentation results with higher accuracy than *BCM3D 1*.*0*. Importantly and in contrast to *BCM3D 1*.*0*, generation of intermediate image representations does not require image deconvolution as a pre-processing step. Deconvolution can lead to noise amplification^31^ that then leads to false positive object segmentation with physiologically unreasonable shapes. We show that, using intermediate image representations, experimentally acquired biofilm images can be successfully segmented using CNNs trained with computationally simulated biofilm images – a feature shared with *BCM3D 1*.*0* that provides the flexibility to segment a wide variety of different cell shapes^23^. The segmentation performance of this new approach, which we term *BCM3D 2*.*0*, is superior to *Omnipose* and *Cellpose 2*.*0*, two recently-developed, state-of-the-art, CNN-based cell segmentation approaches, especially for dense biofilms imaged at low SBRs. The improvements in segmentation accuracy of *BCM3D 2*.*0* enables accurate multi-cell tracking, which is demonstrated using 3D simulated and experimental time-lapse biofilm images.

## Methods

### Lattice Light Sheet Microscope Imaging of Bacterial Biofilms

Fluorescence images of bacterial biofilms were acquired on a home-built lattice light sheet microscope (LLSM). LLSM enables specimen illumination with a thin light sheet derived from a 2D optical lattice^32, 33^; here, an intensity uniform light sheet was produced by dithering a square lattice. The average illumination intensity across the light sheet was less than 1 W/cm^2^. The sub-micrometer thickness of the light sheet is maintained over a propagation distance of ∼30 μm to achieve high resolution, high contrast imaging of 3D specimens comparable to confocal microscopy but with lower concomitant photobleaching and phototoxicity. Widefield fluorescence images of illuminated planes in the specimen are recorded on a sCMOS detector (Hamamatsu ORCA Flash v2). 3D biofilm images were acquired by translating the specimen through the light sheet in 200 nm step sizes using a piezo nano-positioning stage (Mad City Labs, NanoOP100HS). The data acquisition program is written in LabVIEW 2013 (National Instruments).

Kanamycin resistant *S. oneidensis* MR-1, constitutively expressing GFP, were cultured at 30 °C overnight in LB medium with 50 μg/ml Kanamycin. Overnight cultures were diluted 100 times into the same culture medium, grown to an optical density at 600 nm (OD600) of 0.4 – 1.0, and then diluted to OD600 ∼ 0.05 using M9 media with 0.05% (W/V) casamino acids. Poly-l-lysine coated round glass coverslips with the diameter of 5 mm were put into a 24-well plate (Falcon) and 400 μL of diluted cell culture was added to the well. Cells were allowed to settle to the bottom of the well and adhere to the coverslip for 1 hour. After the settling period, the coverslip was gently rinsed with M9 media to flush away unattached cells. Then 400 μL of M9 media (0.05% casamino acids) were added to ensure immersion of the coverslips. The well plate was set in a 30 °C chamber for 72-96 hours to allow dense biofilms to develop. Media were exchanged every 24 hours. Before imaging, the coverslip was rinsed again with fresh M9 media. The rinsed coverslip was then mounted onto a sample holder and placed into the LLSM sample-basin filled with M9 media. GFP was excited using 488 nm light sheet excitation. 3D biofilm stacks were acquired by translating the specimen through the light sheet in 200 nm or 235 nm steps. Each 2D slice was acquired with an exposure time of 5 ms or 10 ms.

Samples for time-lapse images were prepared by the same procedures, except imaging was started after either 24-hour or 48-hour cell attachment period, and the imaging experiment was carried out in LM medium (0.02% (W/V) yeast extract, 0.01% (W/V) peptone, 10 mM HEPES (pH 7.4), 10 mM NaHCO_3_) with a lactate concentration of 0.5 mM.^34^ Time-lapse images were recorded every 30 seconds for 15 minutes or 5 minutes for 5 hours for the two datasets shown in **Figure 5** with the same imaging parameters as detailed above.

### Raw Data Processing

Raw 3D stacks were deskewed and rotated as described previously^35^, but the deconvolution step was omitted. If necessary, background subtraction can be applied to reduce background signal. 3D images were rendered using the 3D Viewer plugin in Fiji^36^ or ChimeraX^37^. Sample drift over the course of a time-lapse imaging experiment was corrected by Correct 3D Drift^38^, a Fiji plug-in that performs registration by phase correlation, a computationally efficient method to determine translational shifts between images at two different time points.

### Generation of simulated biofilm images

Data for CNNs training was computationally generated as described previously^23^. Briefly, CellModeller^39^, an individual-based computational model of biofilm growth, was used to simulate growth and division of individual rod-shaped cells in a population (**Figure 1a**). A minimum distance criterion between cells is imposed at each time point to alleviate cellular collisions that are due to cell growth. We chose cell diameter and cell length (d, l) parameters consistent with the bacterial species under investigation, namely (1 μm, 3 μm) for *E. coli*^*40*^, and (0.6 μm, 2 μm) for *S. oneidensis*^*41*^. Training data should closely represent the experimental data to ensure optimal segmentation results. Unrepresentative cell diameter and cell length parameters can result in over- or under-segmentation errors and the predictions of non-physiological cell shapes (**Figure S1**). 3D fluorescence intensity images (**Figure 1b**) were generated by convolving randomly positioned fluorophores in the cytoplasm or the membranes of simulated cells (**Figure 1cd**) with experimentally measured point spread functions (PSFs), and then adding experimentally measured background and noise (Poisson detection noise, based on the summed background and signal intensities, as well as Gaussian read noise, experimentally calibrated for our detector at 3.04 photons per pixel on average)^42^.

**Figure 1.**
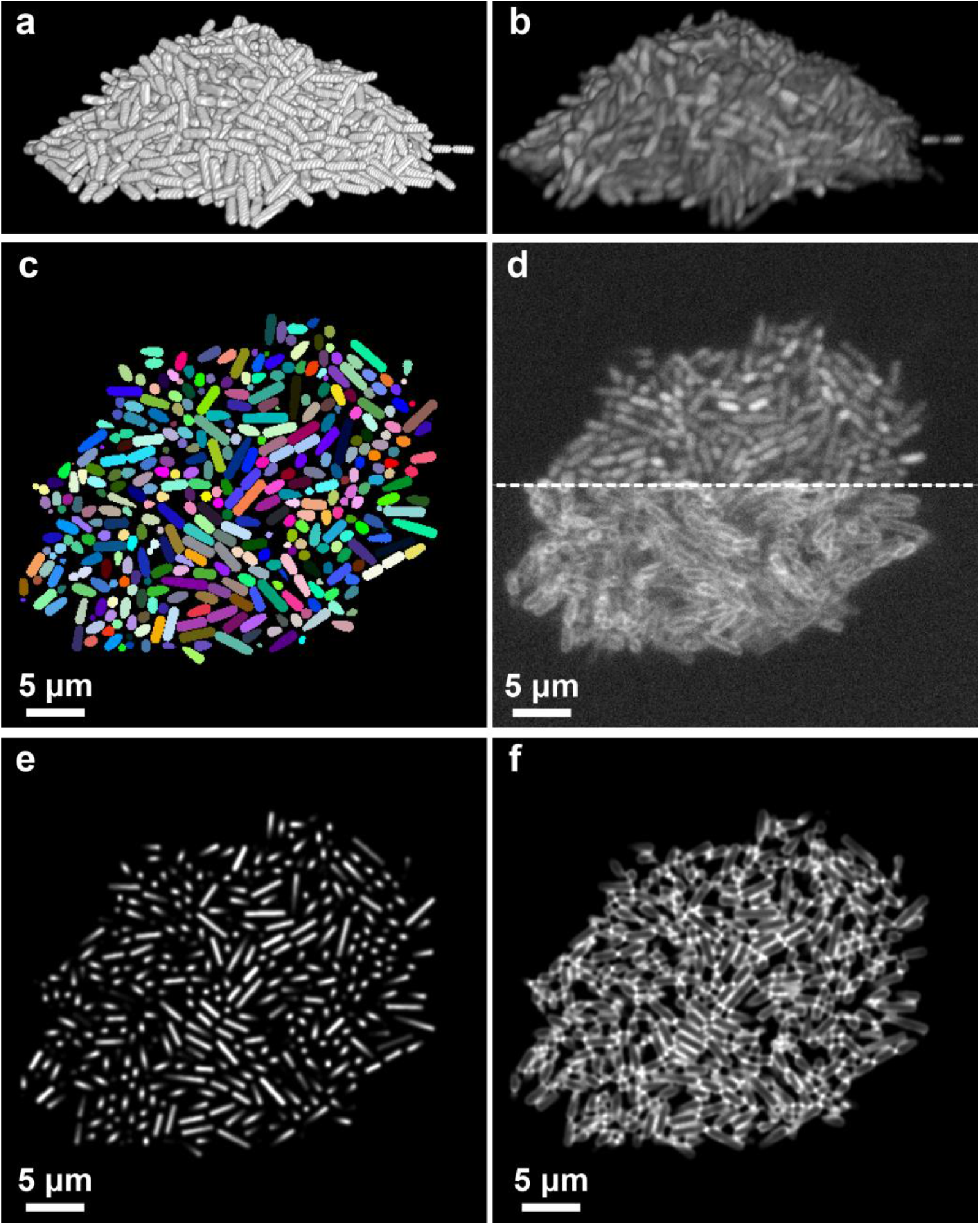
Simulation of fluorescent biofilms images and intermediate image representations. (a) Cell arrangements obtained by CellModeller. (b) Simulated 3D fluorescence image based on the cell arrangements in a. (c) Ground truth information of a 2D slice. Different cells are shown in different colors and intercellular spaces (background voxels) are displayed in black. (d) 2D slices of the simulated fluorescence image corresponding to the ground truth shown in c. The upper panel shows cells containing cytosolic fluorophores, the lower panel shows cells with fluorescently stained membranes. (e and f) Intermediate image representations generated from the ground truth information shown in c. See text for details.

The fluorescence signal intensity in the simulated images was adjusted to match the signal to background ratios (SBRs) of experimentally acquired data. To estimate the SBRs of both simulated and experimental images, we manually selected 10 ‘signal’ and 10 ‘background’ regions in the images using the Oval tool in Fiji and calculated their means respectively. A ‘signal’ region is defined to be any region that contains only pixels within a cell (foreground) and a ‘background’ region contains only pixels outside cells, but in regions that are close to the cells. These regions are judged by the researchers rather than by any computer algorithm to ensure accuracy. The SBR was then calculated by dividing the mean signal intensity by the mean background intensity. Consistent with our previous results (Zhang *et al, Nature Communications*, 2020), the SBR is a good metric quantifying “difficulty to segment” in simulated data, which has homogeneous cells densities and the exact same biofilm architectures (i.e. cell positions). For heterogeneous cell densities in experimental biofilms, the SBR can vary considerably through space. We therefore quantify the SBR in experimental images locally at regions of highest cell density, but we note that this metric should only be used qualitatively and we refrain from making any direct, quantitative comparison of segmentation performance between biofilms of *different* architectures. To estimate the local density of a biofilm, the image was partitioned into several 3D tiles 64 by 64 by 8 voxels in size and the total cell volume contained in each tile was divided by the tile volume. The reported cell density was computed as the average of the 10 densest tiles for each dataset.

### Generation of intermediate image representations

To generate ‘distance to nearest cell exterior’ images (**Figure 1e, Figure S2**) from ground truth data, the Euclidean distance of each voxel inside a cell to the nearest voxel not belonging to that cell was calculated. The so-obtained distances were then normalized to the maximum value of that cell (**Figure S2c**). In order to obtain a steeper gradient in distance values, the distance values were additionally raised to the third power (**Figure S2d**), so that the resulting images show highly peaked intensity near the cell center. In a final step, the ‘distance to nearest cell exterior’ images were smoothed by Gaussian blurring (kernel size = 5 voxels in each dimension) (**Figure S2e**).

To help distinguish touching cells, we calculated a second image representation, the ‘proximity enhanced cell boundary’ image (**Figure 2f, Figure S2**). First, we subtracted the normalized distances to the nearest voxel not belonging to this cell (**Figure S2c**) from the binary map (**Figure S2f**). Second, we calculated the inverse of the Euclidean distance of each voxel inside a cell to the nearest voxel belonging to another cell, an intermediate image representation that has been proven useful to prevent objects merging in 2D^27^ (**Figure S2g**). These two intermediate images were then multiplied together (**Figure S2h**) and small holes in the resulting images (**Figure S2h inset**) were filled using grayscale closing (**Figure S2i inset**). The resulting intermediate images provides a complete boundary of an object but also highlights whether the boundary is in close proximity to any other objects. Compared to previous methods that only provide a complete boundary or only provide boundary areas that are close to any other objects, this new intermediate image representation provides a more informative boundary representation. In a final step, the ‘proximity enhanced cell boundary’ images were smoothed by Gaussian blurring (kernel size = 5 voxels in each dimension) (**Figure S2i**).

**Figure 2.**
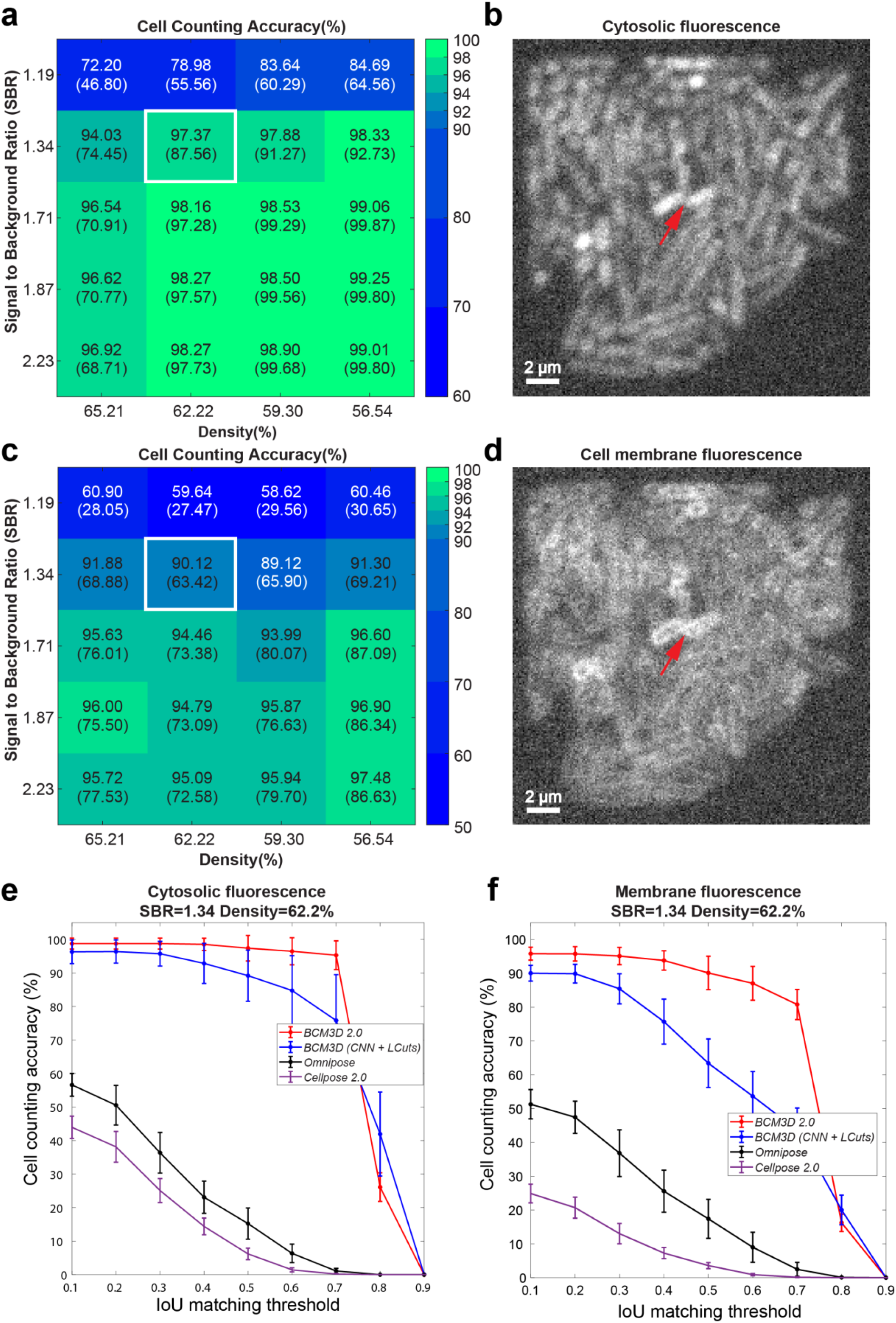
Performance of *BCM3D 2*.*0* on previously unseen simulated biofilm images. **(a)** Cell counting accuracy (using an Intersection-over-Union (IoU) matching threshold of 0.5 for each segmented object) averaged over *N*=10 replicate datasets for cells labeled with cytosolic fluorophores. IoU is well-documented metric^23, 30, 56^ quantifying the amount of overlap between predicted cell and actual cell volumes. **(b)** Example image of cells labeled with cytosolic fluorophores (Cell density = 62.2%, SBR = 1.34, indicated by white rectangle in panel a). **(c)** Cell counting accuracy (using an IoU matching threshold of 0.5 for each segmented object) averaged over *N*=10 replicate datasets for cells labeled with membrane-localized fluorophores. **(d)** Example image of cells labeled with membrane-localized fluorophores (Cell density = 62.2%, SBR = 1.34, indicated by white rectangles in panel c). The red arrow indicates a close cell-to-cell contact. **(e** and **f)** Comparison of segmentation accuracies achieved by *BCM3D 1*.*0 (*CNN *+ LCuts), Omnipose* trained from scratch, *Cellpose 2* fine-tuned and *BCM3D 2*.*0* for cytosolic and membrane labeling, respectively (SBR = 1.34, cell density = 62.2%). Segmentation accuracy is parameterized in terms of cell counting accuracy (*y* axis) and IoU matching threshold (*x* axis). Each data point is the average of *N*=10 independent biofilm images. Data are presented as mean values ± one standard deviation.

### Training the convolutional neural network

To generate the above-mentioned intermediate image representations from experimental data, we trained 3D U-Net-based CNNs with residual blocks using the CSBDeep Python package^19^. Residual blocks allow the model to internally predict the residual with regard to inputs for each layer during training. This strategy provides better performance, because solvers are more efficient in solving residual functions than unreferenced functions, and it helps alleviate vanishing or exploding gradients problems for deep neural networks^43^. We employed a network architecture depth of 2, a convolution kernel size of 3, 32 initial feature maps, and a linear activation function in the last layer. Increasing U-Net depth or numbers of initial features didn’t produce superior results for our test cases (**Figure S5**). To achieve robust performance, we trained this network using ten to twenty simulated biofilm images with randomly selected cell densities and signal-to-background ratio. To ensure the broad applicability of these networks, half of these images were biofilms containing cells expressing cytosolic fluorescence and the other half were biofilms containing membrane-stained cells (see **Figure 1d**). The loss function was taken as the mean absolute error (MAE) between the generated and the target images. The networks were trained for 100 epochs with 100 parameter update steps per epoch and an initial learning rate 0.0004 (**Figure S6**). The learning rate is reduced by half if the validation loss is not decreasing over 10 epochs. We selected the best weights based on performances on validation set for all the processing steps described below. Using these parameters, it took approximately 1 hour to train the CNNs on a NVIDIA Tesla V100 GPU with 32 GB memory.

To obtain instance segmentation results from intermediate image representations predicted by trained CNNs, we applied single- and multilevel Otsu thresholding^29, 44^, and seeded watershed^28^ (scikit-image Python library^45^, **Figure S3** and **S4**). The reader is referred to the **Supplemental Information** for detailed, step-by-step explanations.

To test whether segmentation objects have physiologically reasonable cell shapes, we separately trained a 3D CNNs based classification model using tensorflow 2.0. We adapted a network architecture from Zunair et.al.,^46^; mainly includes three 3D convolutional layers, one global average pooling layer and a sigmoid activation function in the last layer. To achieve robust performance, we trained this network using 733 manually confirmed segmentation objects from experimental data (411 reasonable shaped objects, 322 oddly shaped objects). Training data were augmented by rotation and flip. The loss function was taken as the binary cross entropy between the model output and the corresponding target value. The networks were trained for 100 epochs with a batch size of 5 and an initial learning rate 0.0002. The learning rate is reduced by a half if the validation loss is not decreasing over 15 epochs. Using these parameters, it took approximately 17 mins to train the CNNs on a NVIDIA Tesla V100 GPU with 32 GB memory.

### Tracking

*Simpletracker* in MATLAB was used to build tracking graphs and spatially resolved lineage trees^47^. *Simpletracker* implements the Hungarian algorithm and nearest neighbor trackers for particle tracking that links particles between frames in 2D or 3D. We used 1 μm and 1.5 μm as the maximum distance threshold for cell linking for simulated and experimental data, respectively. We used the nearest neighbor algorithm to associate the centroids of segmented objects in subsequent frames, such that the closer pairs of centroids are linked first. In order to determine a cell division event, a distance threshold of 1 μm and 1.5 μm for simulated and experimental data, respectively, a cell volume threshold of 1.5 (parent cell should be 1.5 times larger than the daughter cell), and a cell length threshold of 1.5 (parent cell should be 1.5 times longer than the daughter cell), were used to determine parent-daughter relationships between cell pairs on consecutive frames.

### Performance evaluation

Segmentation accuracy was quantified as cell counting accuracy and cell shape estimation accuracy. The cell counting accuracy (CA) was calculated as previously described^23^:

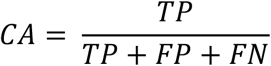

where, TP is the number of truth positive objects, FP is the number of false positive objects, and FN is the number of false positive objects. Cell shape estimation is evaluated by two separate measures. Single-cell segmentation accuracy (SSA) takes the mean Intersection-over-Union (IoU) value (aka the Jaccard index^48^) over segments that have a matching ground truth/manual annotation object:

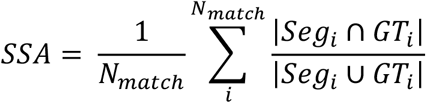

where, |*Seg*_*i*_ ∩ *GT*_*i*_| is volume of overlap between the predicted object and the ground truth object, and |*Seg*_*i*_ ∪ *GT*_*i*_| is the volume enclosed by both the predicted object and the ground-truth object. We note that the SSA metric can take on high values even if the shape of a segmented object does not accurately represent the shape of the corresponding ground truth object. For example, a predicted round object with a diameter of 20 covered by a ground truth square object with a length of 20 gives a 0.8 IoU value, which could be interpreted as good performance. From a biological perspective however, this would signify a substantial inaccuracy in shape estimation. To measure differences in cell shape in a more discriminating way, we additionally computed a single-cell boundary F1 score (SBF1)^49^. The SBF1 of the abovementioned square vs circular object example is 0.67. The SBF1 score is computed as

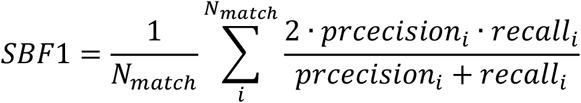

where precision is the ratio of matching boundary points in a matched segmentation object to the total points of its boundary. Similarly, recall is the ratio of the matching boundary points to the total points of ground truth boundary. According to the definition of boundary F1 score^50^, a distance error tolerance is used to decide whether a point on the predicted boundary has a match on the ground truth boundary. For our 3D data, we use √3 voxels.

To quantify tracking accuracy, we used the acyclic oriented graph metric (*AOGM*)^51^. The *AOGM* value is calculated as the weighted sum of the number of graph operations required to convert the estimated graph to the ground truth graph, i.e.:

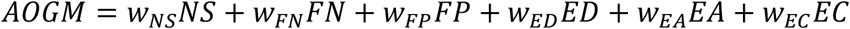

The tracking accuracy can then be computed using a normalized *AOGM* value, where *AOGM*_*0*_ is the number of operations to build the ground truth graph from an empty graph:

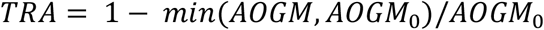

There are three types of graph operations that are associated with detection errors: the number of false negatives (*FN*), the number of false positives (*FP*), and the number of missed splits (*NS*: *m* reference cells (*m* > 1) are assigned to a single segmented cell); and three types of graph operations that are associated with object linking: edge deletion (ED), addition (*EA*), and alteration of the semantics of an edge (*EC*: The semantics of an edge can either represent the same cells over time or represent a parent-daughter relationship). To focus on object matching over time (i.e. the association performance of the algorithm), we used an equally weighted sum of the lowest number of graph operations on edges only (*TRA_edge*). To give a more comprehensive view, we used an equally weighted sum of the number of graph operations on all six operations (*TRA_full*).

To estimate tracking accuracy for experimental data, we manually traced a small subset (n = 25) ancestor cells over time based on *BCM3D 2*.*0* segmentation masks. Two researchers performed tracking independently, manually determining parent-daughter relationships within the lineages originating from the ancestor cells. This lineage information was then used to compute *TRA_edge*.

## Results

### Cell segmentation using intermediate image representations

High cell density and low SBR datasets are encountered often in biofilm research, especially when cells touch each other and biofilms extend farther into the vertical (*z*-) dimension, so that light scattering becomes a pronounced background contribution^52^. We therefore sought to specifically improve bacterial cell segmentation accuracy for high cell density and low SBR biofilm images. Our previous approach (*BCM3D 1*.*0*) relied on deconvolution as a preprocessing step to sharpen the image and to increase the SBR. However, deconvolution can introduce artifacts into an image, such as ringing^53^, and noise amplification^54^, and thereby introduce errors into the segmentation results. The segmentation pipeline of *BCM3D 2*.*0*, in contrast, works on the raw image data directly without the need for deconvolution.

We compared two commonly used cell labeling approaches, namely cell interior labeling through expression of cytosolic fluorescent proteins and cell membrane staining with membrane-embedded fluorescent dyes. For cell interior labeling (**Figure 2ab**), *BCM3D 2*.*0* consistently produces cell counting accuracies of >95% for SBRs > 1.3 and cell densities < 65%. A clear drop-off in cell counting accuracy is observed for SBRs of 1.19 but cell counting accuracies of >70% are still achieved even for high cell densities of 65%. Importantly, the performance of *BCM3D 2*.*0* on low SBR datasets represents a substantial improvement (>20%) over the performance of *BCM3D 1*.*0*. Membrane staining (**Figure 2cd**) produces even more challenging images for segmentation, due to the less pronounced fluorescence intensity minima between cells (red arrow in **Figure 2bd**). We again observe a drop in cell counting accuracy for SBRs of 1.19. This drop-off is however much less pronounced than for the previous results obtained with *BCM3D 1*.*0*, and represents an even larger (>29%) improvement over *BCM3D 1*.*0* for such extremely low SBR datasets. Visual inspection of slices through the image volumes (**Figure 2bd**) reveals that even for SBR = 1.3, the cell bodies are difficult to distinguish for expert human annotators, especially for membrane-stained cells. Despite the low contrast in the SBR = 1.3 datasets, *BCM3D 2*.*0* is still able to achieve >90% cell counting accuracies, which, depending on cell density, represents a 6-26% increase over the performance of *BCM3D 1*.*0*.

To determine the improvement in cell shape estimation, we evaluated the cell counting accuracies as a function of IoU matching threshold for a SBR of 1.3 and a cell density of 62% (the IoU matching threshold is a quantitative measure of cell shape similarity relative to the ground truth). The cell counting accuracies obtained by *BCM3D 2*.*0* are consistently higher than *BCM3D 1*.*0* (CNN + *LCuts*) and substantially higher than *Omnipose* and *Cellpose 2*, especially for IoU matching thresholds larger than 0.5, indicating that cell shapes in this high density, low SBR dataset are most accurately estimated by *BCM3D 2*.*0* (**Figure 2ef**). A similar trend is observed for single-cell segmentation accuracy and single-cell boundary F1 score^49^ – two additional metrics for segmentation accuracy (**Table 1**). We note that we trained *Omnipose, Cellpose 2*, and *BCM3D 2*.*0* using the same simulated training data for this comparison. Consistent with previous findings^55^, *Cellpose*, the precursor algorithm to *Omnipose*, did not produce physiologically-reasonable cell shapes (**Figure S7**). Taken together, these results establish that more robust cell segmentation can be achieved using the *BCM3D 2*.0 image processing pipeline, which uses CNNs to generate two distinct intermediate image representations for subsequent mathematical processing.

**Table 1.** Quantitative comparison of single cell level segmentation accuracy between *BCM3D 1*.*0* and *BCM3D 2*.*0*.

|  |  | Cytosolic labeling |  | Membrane labeling |  |
| --- | --- | --- | --- | --- | --- |
|  |  | SSA* | SBF1** | SSA* | SBF1** |
| *BCM3D 1.0* | (CNN + *LCuts*) | **0.796 ± 0.021** | 0.983 ± 0.008 | 0.756 ± 0.009 | 0.961 ± 0.007 |
| *BCM3D 2.0* |  | **0.791 ± 0.004** | **0.995 ± 0.001** | **0.773 ± 0.005** | **0.988 ± 0.002** |
| *Omnipose* | | 0.599 ± 0.011 | 0.868 ± 0.017 | 0.615 ± 0.021 | 0.824 ± 0.025 |
| *Cellpose 2* | | 0.566 ± 0.016 | 0.825 ± 0.016 | 0.572 ± 0.008 | 0.820 ± 0.009 |

\*SSA and \*\*SBF1 estimate how accurately the shape of a matched object compare with it of the corresponding ground truth. Here, the IoU threshold is 0.5 and the distance error tolerance for
SBF1 is $\sqrt{3}$ voxels. (See Methods for details). Data are presented as mean values ± one standard deviation, the best performance (if different within error) is marked in bold.

### Segmentation of experimentally obtained biofilm images

To test the performance of *BCM3D 2*.*0* on experimental data, we first tested *BCM3D 2*.*0* on a previously published *E*.*coli* biofilm image, for which manual annotation masks are available ^23^. For this dataset, which features a relatively low cell density, *BCM3D 2*.*0* performed on par with *BCM3D 1*.*0* (CNN + LCuts), but again outperformed *Omnipose* and *Cellpose 2* (**Figure S8)**. (We only considered recently developed 3D instance segmentation approaches for these comparisons. In previous work^23^, we established that *BCM3D 1*.*0* (CNN + *LCuts*) outperformed both *Cellpose* and the segmentation algorithm developed by Hartmann *et al*.) To further demonstrate the versatility of *BCM3D 2*.*0*, we acquired images of a thick *S. oneidensis* biofilm expressing GFP that has an order of magnitude more cells. Visual inspection of the segmentation results obtained by applying *BCM3D 2*.*0*, showed physiologically reasonable cell shapes for a majority of segmented objects (**Figure 3**). For this biofilm image, which has a good SBR, *BCM3D 2*.*0* outperforms all other approaches, including *Omnipose, BCM3D 1*.*0* (CNN + *LCuts*) and *Cellpose 2*.*0* (**Figure S9)**.

**Figure 3.**
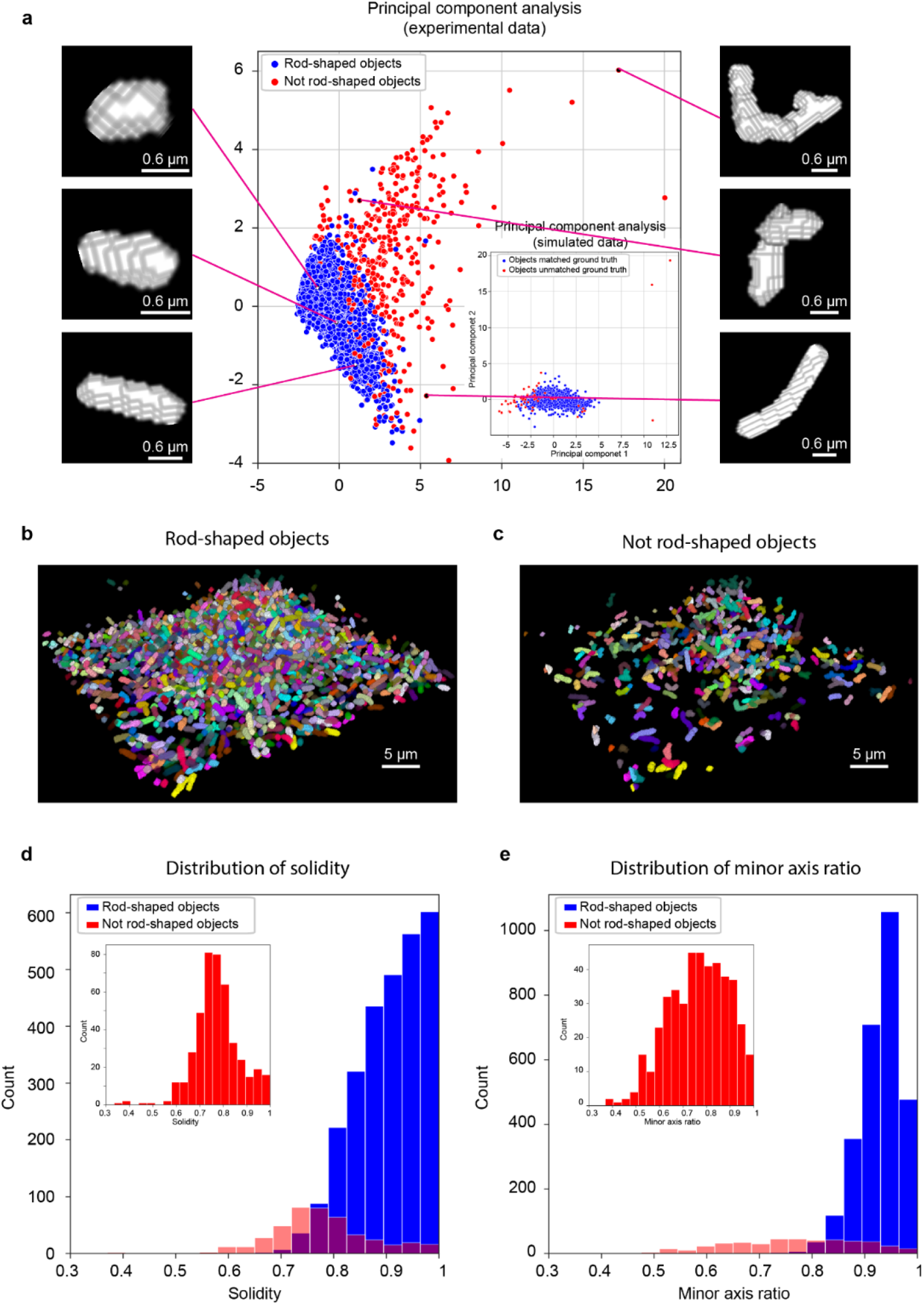
Performance of *BCM3D 2*.*0* on experimental biofilm images. (a) Principal component analysis of the segmentation objects (obtained from experimentally acquired images) that were classified by a pre-trained 3D CNN as either physiologically reasonable rod-shaped cells or oddly shaped (not-rod shaped) cells. Three examples cell shapes of each class are shown to the right and left, respectively. Inset: same analysis on simulated biofilm images. (b) 3D segmentation results for objects classified as physiologically reasonable rod-shaped cells. (c) 3D segmentation results for objects classified as oddly shaped. (d) and (e) Comparison of the solidity and minor axis ratio distributions of rod-shaped and oddly shaped objects.

The above quantitative comparisons were made with reference to manual annotation results made in selected 2D slices. Manual annotation of 3D biofilm images is however too time consuming for thousands of cells distributed in 3D space^23^. We therefore chose to additionally assess the segmentation accuracy using representative morphological observables that are available after segmentation, namely object volume, object solidity (volume fraction of the object as compared to the smallest convex polygon that encloses it), major axis length, longer minor axis length, and the ratio of the two minor axes lengths (longer minor axis divided by the shorter one). We performed principal component analysis (PCA) using these morphological observables and project each segmented object onto a plane spanned by the first two principal components. For simulated data (for which the ground truth is known) this approach shows a distribution for which the correctly segmented objects are concentrated near the origin, whereas the incorrectly segmented objects are predominantly located at the periphery (**Figure 3a** inset).

We applied the same PCA approach to experimental segmentation results obtained for a *S. oneidensis* biofilm containing ∼3000 cells. Similar to simulated data, most of the segmentation objects cluster near the origin of the two principal component axes (**Figure 3a**). However, several segmented objects are asymmetrically scattered around the periphery of the distribution. Inspecting the 3D shapes of a few manually selected objects revealed that, consistent with simulated data, physiologically reasonable cell shapes cluster near the center of the distribution, while oddly shaped objects predominantly localize at the periphery. To automatically separate oddly shaped objects from the physiologically reasonable, rod-shape shaped objects, we trained a 3D CNN (independent of *BCM3D*) with manually validated segmentation objects (obtained from experimental data, see methods). The trained network efficiently (accuracy of 97% on the validation set) separates rod-shaped objects (∼82% of total) from oddly shaped objects (∼18% of total). This classification enables the display of both subpopulations separately even though they are completely intermixed in 3D space (**Figure 3bc**). In contrast, *Omnipose* and *Cellpose 2*.*0* have 40% and 5% rod-shaped objects respectively identified by the shape classifier on this dataset.

We next compared the distributions of solidity and minor axis ratio between rod-shaped and oddly shaped populations. Rod-shaped objects are characterized high values of solidity and minor axis ratio (**Figure 3de**). In contrast, solidity and minor axis ratio for oddly shaped objects take on values less than one and thus show a much broader distribution (**Figure 3de** insets). These results show that, when using *BCM3D 2*.*0*, ∼82% of cells are segmented with physiologically reasonable cell shapes. The remaining 18% of cells can then be subjected to further processing to identify and correct the remaining segmentation errors^23, 24, 49^ and/or be subjected to further scrutiny to determine whether they are due to aberrant cell shapes exhibited by sick or intoxicated cells^55^.

### Accurate *BCM3D 2*.*0* segmentation enables multi-cell tracking in biofilms

Simultaneous multi-cell tracking and lineage tracing is critical for analyzing single-cell behaviors in bacterial biofilms. We asked whether the cell segmentation performance of *BCM3D 2*.*0* was sufficient to enable accurate tracking of individual cells in biofilms. To address this question, we employed a tracking-by-detection approached using simulated biofilm images of different SBRs (**Figure 4a**). We evaluated tracking accuracy, as a function of SBR, using the widely used *TRA* metrics based on Acyclic Oriented Graph Matching (AOGM)^51^. In acyclic oriented graphs, cells in different time frame are represented as vertices and linkages between cells from frame-to-frame are represented as edges. When the cells (vertices) are placed at their actual (x,y,z) spatial coordinates, then the cell linkages (edges) represent the branches of a spatially resolved lineage tree (**Figure 4b**). The *TRA* metrics quantify the minimum number of elementary graph operations that are needed to transform an estimated graph into a ground truth graph. *TRA_edge* considers three edge operations only, while *TRA_full* considers all six graph operations^51^.

**Figure 4.**
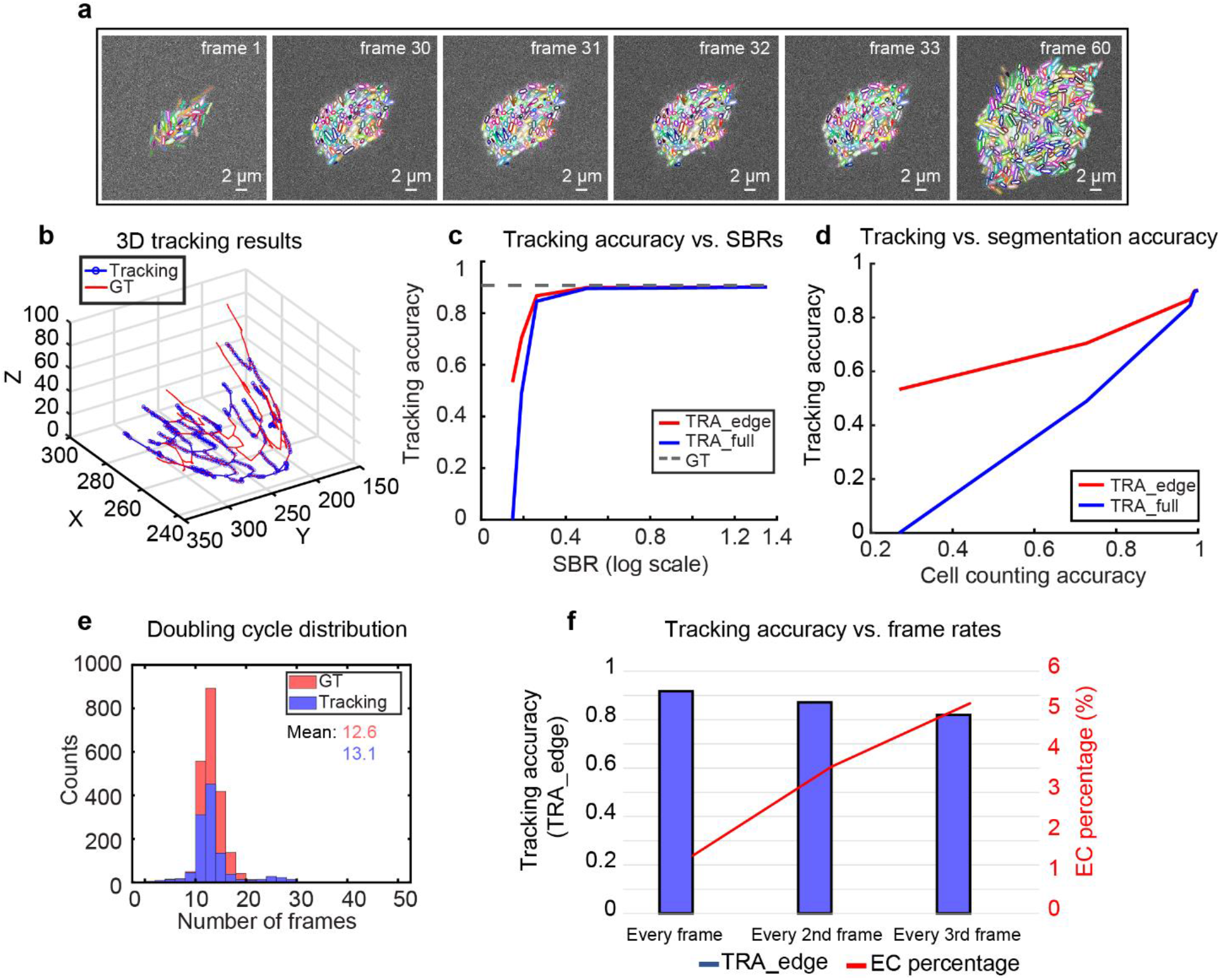
Multi-cell tracking in simulated biofilms. (a) Simulated fluorescence time-lapse images of growing *E. coli*-like biofilm. The SBRs of these images are ∼1.65. Contours are color-coded based on segmentation and tracking results. (b) An example of 3D tracking and lineage tracing for simulated biofilm images. For clarity, spatial trajectories and lineages originating from only a single ancestor cell is displayed. The estimated graph is shown in blue and the corresponding ground truth graph is shown in red. The entire biofilm contains over sixty graphs of this type. (c) Two AOGM metrics calculated as *TRA_edge* and *TRA_full* are plotted against image SBR. The grey dashed line indicates tracking of the GT segmentation using the same tracking algorithm. (d) Same data as in panel c plotted as a function of cell counting accuracy at IoU = 0.5, a segmentation accuracy metric that increases for increasing SBR in the raw images^23^. (e) Doubling cycle distribution of simulated data and corresponding tracking results. A completed cell cycle is defined as a track in which the parent cell is able to split twice. This threshold results in a lower count numbers of estimated cell division, but does not alter the shape of the distribution. (f) *TRA_edge* (left axis) and edge correction (EC) percentage (right axis) for different temporal sampling rates. EC percentage indicates how many parent-daughter relationships are misassigned based on the tracking results.

To link the same cells across two different time points, we used a nearest neighbor algorithm^57^. When using spatial distance as the sole metric for cell linking, the AOGM tracking accuracy has a positive correlation with SBR (**Figure 4c**), which highlights the importance of accurate cell segmentation in multi-object tracking-by-detection^58^. *BCM3D 2*.*0* enables a tracking accuracy that is similar to the ground truth tracking accuracy (same nearest neighbor tracking algorithm applied to the ground truth segmentation masks) for SBRs of 1.65 and higher. We note that, given the high cell density in this test dataset, the ground truth tracking accuracy does not reach the optimum (100%) even with error-free segmentation. This is due to inherent limitations in how mother daughter relationships are assigned. At SBR’s less than 1.65, tracking accuracy decreases rapidly due to the lack of consistent segmentation results. The importance of accurate segmentation is clearly evidenced by the linear dependence of *TRA* as a function of cell counting accuracy (**Figure 4d**).

Another key factor for simultaneous multi-object tracking is the time resolution^58^. The relative movement (RM) of objects from frame to frame is therefore a useful metric to quantify the level of difficulty for cell tracking. The relative movement (*RM*_*i,j*_) in time frame *i*, for a given cell *j* is defined as the ratio between the distance of cell *j* to itself between frame i and *i*+1 and the distance of cell *j* in frame *i* to its closest neighbor at frame *i* + 1. The <*RM*> metric is then the average *RM*_*i,j*_ of all cells for each frame^59^. A dataset with <*RM*> values of 1 or more means that any tracking method that considers only distance (and distance related features) is likely to fail, whereas a dataset with a <*RM*> value of less than 0.5 is considered challenging^59^. For the simulated biofilm images here, *RM*∼0.2, which indicates that the time resolution may be good enough for single cell tracking using a nearest neighbor algorithm. Indeed, under these conditions, many cells can be tracked for several generations (**Figure 4b**). However, even at *RM*∼0.2, some cell division events are missed, so that a few branches of the lineage tree are not successfully traced. Even so, the subset of correctly detected cell division events allows for the estimation single-cell doubling cycles in the biofilm (**Figure 4e**). To quantify these trends, we tested how time resolution affects tracking accuracy. When the time resolution is decreased by a factor of two and three, the *TRA_edge* metrics decrease from 91% to 87% and 81%, respectively. The percentage of the parent-daughter misassignment error, quantified as the edge-correction (EC) error over the number of total errors, increases from 1.4% to 3.6 and to 5.2 % (**Figure 4f**). Taken together, these results show that segmentation based multi-object tracking accuracy is highly dependent on segmentation accuracy (which depends on image SBR and cell density^23^), as well as time resolution. It is critical to consider these parameters, when single-cell resolved observables, such as cell trajectories, single cell volume increases, and single-cell doubling times, need to be measured.

### Multi-cell tracking in the initial phase of *S. oneidensis* biofilm

Cell segmentation and subsequent multi-cell tracking in experimentally acquired 3D images presents additional challenges that were not modeled in the computationally simulated data. These challenges include optical aberrations in the imaging system, broader cell shape distributions in experimental biofilms, cell motility, and association and dissociation dynamics of individual cells to and from the biofilm. To determine whether the *BCM3D 2*.*0* segmentation results enable improved multi-cell tracking using a nearest neighbor algorithm, we manually traced a subset of ancestor cells over the course of a 15-minute 3D biofilm movie acquired with a time resolution of 30 seconds (**Figure 5ab**). Manual determination of cell-to-cell correspondences in consecutive image volumes generated 583 cell-cell and 3 parent-daughter linkages. Taking this manual annotation as the reference graph, the *RM* metric was determined to be ∼0.2 and the *TRA_edge* metric was determined to be 93.5%. Steadily increasing single cell volumes for four selected cells allowed us to measure growth rates of 7.4×10^−3^, 3.8×10^−3^, 3.4×10^−3^, and 0.6×10^−3^ μm^3^/min (**Figure 5c**). Cell division events are also readily detected by the algorithm as a sudden decrease in cell volume. In two of the four selected cases, cell division led to the dispersal of the daughter cell. We found a high number of cell dispersion events resulting in the termination of trajectories, most often right after cell division (**Figure 5c**).

**Figure 5.**
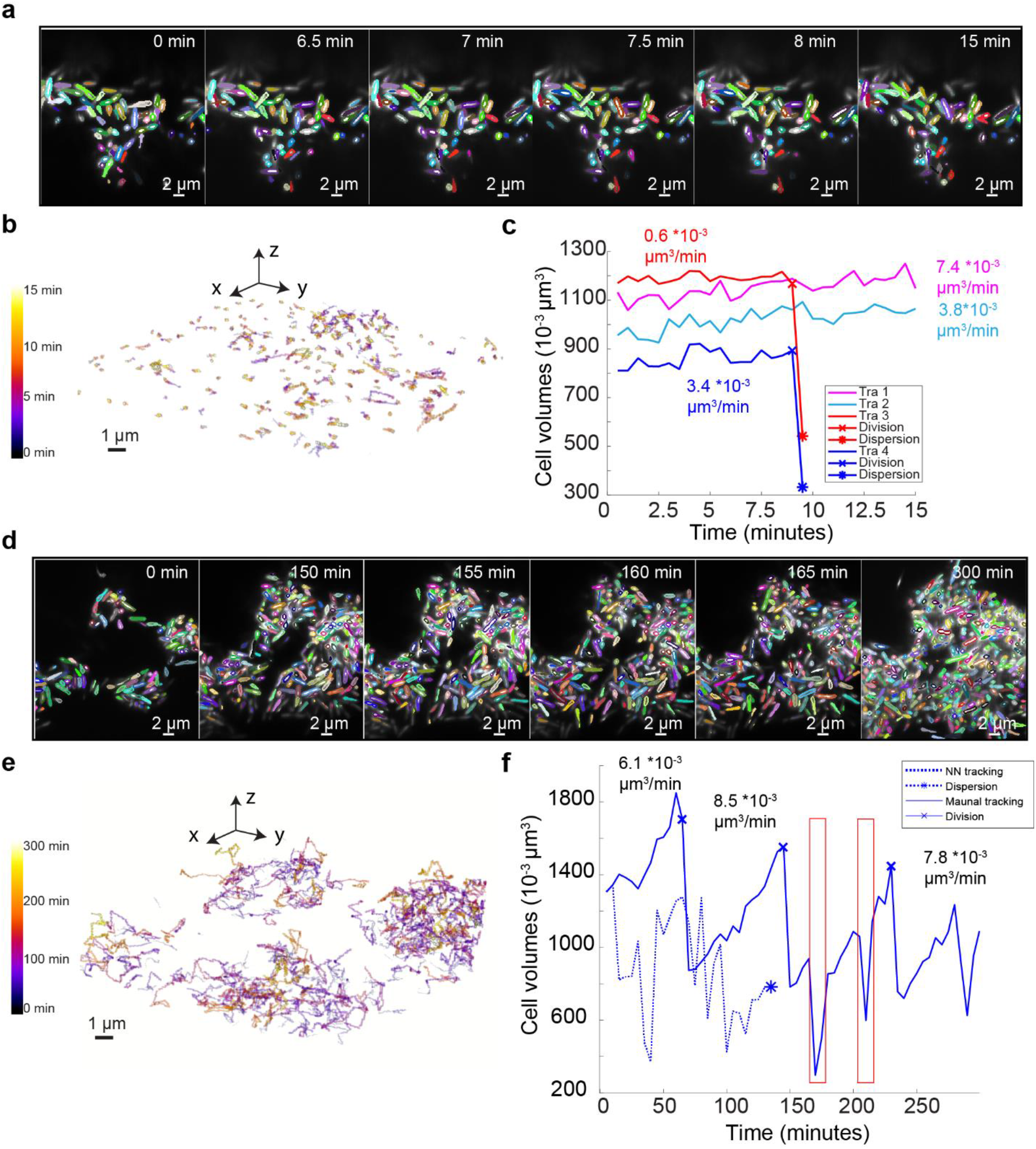
(a) Experimentally acquired fluorescence time-lapse images of a growing *S. oneidensis* biofilm with overlaid single-cell segmentation contours. Images were acquired every 30 seconds for 15 minutes. Corresponding cells in different frames are displayed in the same color. (b) Individual cell trajectories in the biofilm shown in panel a. Cells move very little during the short 15-minute imaging time. (c) Cell volumes over time for four example cells. Cell division and dispersion events are indicated for each trajectory. Single-cell growth rate was measured by calculating the slope of linearly fitted line for each curve (d) Experimentally acquired fluorescence time-lapse images of a growing *S. oneidensis* biofilm with overlaid single-cell segmentation contours. Images were acquired every five minutes for five hours. Corresponding cells in different frames are displayed in the same color. (e) Individual cell trajectories in the biofilm shown in panel a. Cell displacements are more pronounced over the 5-hour imaging time. (f) Evolution of cell volumes over time for a single selected cell. The solid line represents a manually annotated trajectory, and the dashed line represents the nearest neighbor tracking trajectory of the same initial cell. Cell division and dispersion events are indicated on each trajectory. The single-cell growth rate was measured by calculating the slope of the trajectory segment between consecutive cell division events. The red arrow and the boxes indicate periodic underestimation of the cell volume due to oversegmentation.

Although *BCM3D 2*.*0* in combination with high-frame rate imaging enables accurate cell tracking, it may not be feasible to maintain high-frame rate volumetric imaging for extended periods of time due to phototoxicity and photobleaching concerns. To further test the limits of nearest neighbor tracking, we tracked *S. oneidensis* biofilm growth for five hours at a time resolution of 5 minutes (**Figure 5de**). During this time period, the number of cells increases from ∼300 to ∼1400 cells. The relative cell motion in this dataset, estimated by the distances between manually tracked cell centroids, is 0.5 ± 0.2 μm (mean ± standard deviation, *N* = 5). The average spacing of the biofilm, as calculated by the average distance of each cell to the nearest neighbor, changes over time. The average spacing for the first frame, estimated by the average distance to the nearest neighbor for each cell, is 1.2 ± 0.6 μm (mean ± standard deviation, *N* ∼ 300), and is 1.1 ± 0.2 μm (mean ± standard deviation, *N* ∼ 1400) for the last frame. We manually traced a subset of founder cells over the course of the experiment, generating 262 cell-cell and 17 parent-daughter linkages. For this manually selected subset, the *RM* metric was ∼0.4 and *TRA_edge* metric was determined to be 80.0%. While the nearest neighbor tracking algorithm is capable of making overall accurate cell-cell linkages for a few consecutive frames, automated nearest-neighbor tracking of the same cells for long time periods and correctly detecting all cell-division events is not readily possible (**Figure 5f**). It is, however, possible for human annotators to track individual cells from the segmentation results under such imaging conditions. Single-cell growth rates and single-cell division times can then be readily extracted (**Figure 5f, Figure S10, Movie S1**). The measured growth rates are in excellent agreement with the values obtained with high time resolution imaging (**Figure 5c**). A small number of segmentation errors can be detected by manual tracking, as indicated by the boxes in **Figure 5f**, but these errors don’t preclude estimations of single-cell observables. These results indicate that quantitate information about single-cell behaviors is contained even in low time resolution 3D movies of bacterial biofilms. Future work will need to focus on extracting the information as accurately as, but faster than, a human annotator.

## Discussion

We expanded the *BCM3D* workflow with a complementary CNN-based processing pipeline, named *BCM3D 2*.*0*, which transfers raw 3D fluorescence images to intermediate image representations that are more amenable to conventional mathematical image processing (specifically, seeded watershed and single- and multi-level Otsu thresholding). Using the *BCM3D 2*.*0* image processing pipeline, unprecedented segmentation results are obtained, especially for challenging datasets characterized by low SBRs and high cell densities. *BCM3D 2*.*0* consistently achieves better segmentation accuracy than *Cellpose 2*.*0* and *Omnipose*, as well as our predecessor algorithm, *BCM3D 1*.*0*, which represented the previous state-of-the-art for 3D cell segmentation in bacterial biofilms.

We used the segmentation results provided by *BCM3D 2*.*0* as the input to a nearest neighbor tracking algorithm to explore the possibility of simultaneous multi-cell tracking in 3D biofilms. We found that accurate, automated multi-cell tracking in 3D time-lapse movies is possible with a nearest neighbor tracking algorithm, if the relative cell movement (RM) between consecutive frames is small. Depending on the type of biofilm and the bacterial species, small RM values can be achieved using moderate time resolutions of 1-5 minutes. However, for the motile *S. oneidensis* cells imaged here, a time resolution of 5 minutes was insufficient for automated nearest-neighbor cell tracking in dense biofilm regions. Tracking accuracy is reduced especially if cells undergo large and unpredictable displacements within the biofilm, and when cells associate or dissociate to and from the biofilm. Even so, single-cell observables, such as growth rates and cell division times can still be extracted based on manual tracking establishing that such information is in fact contained in movies acquired with a time resolution of 5 minutes. Because manual cell tracking is not feasible for biofilms containing thousands of cells, future work will have to focus on extracting this information in an automated manner, efficiently and accurately. Machine-learning based solutions will likely prove to be useful in this context.

A clear experimental solution would be to image biofilms at high time resolutions. However, every fluorescence imaging modality is subject to trade-offs between the achievable spatial and temporal resolution, image contrast (SBR), and phototoxicity and photodamage. If reducing the total radiation dose delivered to the cells is an experimental necessity, light sheet-based microscopy approaches offer substantial advantages over confocal microscopy^52^.

While *BCM3D 2*.*0* is capable of segmenting biofilm datasets of lower SBR than previous methods, further modifications to the image processing pipeline may be needed to enable the tracking of extremely light sensitive or highly motile bacterial species. Additional modifications could be made to further improve segmentation accuracy for datasets with even lower SBRs than those successfully segmented here. On the other hand, more sophisticated tracking algorithms could be employed that consider additional features beyond the Euclidian distances between objects. Recently developed deep learning-based cell trackers for both 2D and 3D data^60, 61^ are primarily designed for mammalian cells with unique cell shapes. These approaches utilize additional similarity features that inform cell linking across different frames. To what extent such approaches would improve tracking of bacterial cells that have more homogeneous cell shapes remains to be explored. Further benefits may also be gained by utilizing punctate cell labeling schemes^15^ or adaptive microscopy approaches, in which higher illumination intensity frames are interspersed with lower illumination intensity frames and the segmentation results in lower SBR frames are informed by the more accurate results obtained in the higher SBR frames.

In summary, the ability to accurately identify and track individual cells in dense 3D biofilms over long periods of time will require the combination of non-invasive fluorescence microscopy approaches for long-term time-lapse imaging and sophisticated image analysis and multi-object tracking tools that provide robust results even for datasets with limited spatial and temporal resolution, and SBR. Here, we have presented an image processing pipeline that enables improved segmentation of dense biofilm-dwelling cells based on 3D fluorescence images of low SBR. The feasibility of simultaneous, automated multi-cell tracking using a simple nearest neighbor tracking algorithm was demonstrated on high time resolution datasets, while manual tracking was possible on lower time resolution datasets. The tools developed here can thus be leveraged to improve our understanding of how coordinated behaviors among biofilm-dwelling cells eventually produce in the macroscopic properties of bacterial biofilms.

## Author Contributions

J. Z., Y. W., and A. G. designed research;

J. Z., Y. W., E. D., T. T., and M. M. performed research;

J. Z., Y. W., E. D., T. T., M. M., S. A. and A. G. analyzed data;

J. Z., Y. W., E. D., T. T., M. M., S. A. and A. G. wrote the paper.

## Acknowledgements

This work was supported in part by the US National Institute of General Medical Sciences Grant 1R01GM139002 (A.G.) and by a grants from the Trans University Microbiome Initiative (TUMI) at the University of Virginia.

## Data Availability

Ground truth, raw images and segmentation results generated in this study are available at Open Science Framework (OSF) (https://osf.io/m4637/). All data used for generating other results presented in this paper are available upon request from the corresponding author.

## Code availability

The code for running all modules is available in a public repository on GitHub: https://github.com/GahlmannLab/BCM3D-2.0

