## Supplemental information for "*BCM3D 2.0*: Accurate segmentation of single bacterial cells in dense biofilms using computationally generated intermediate image representations"

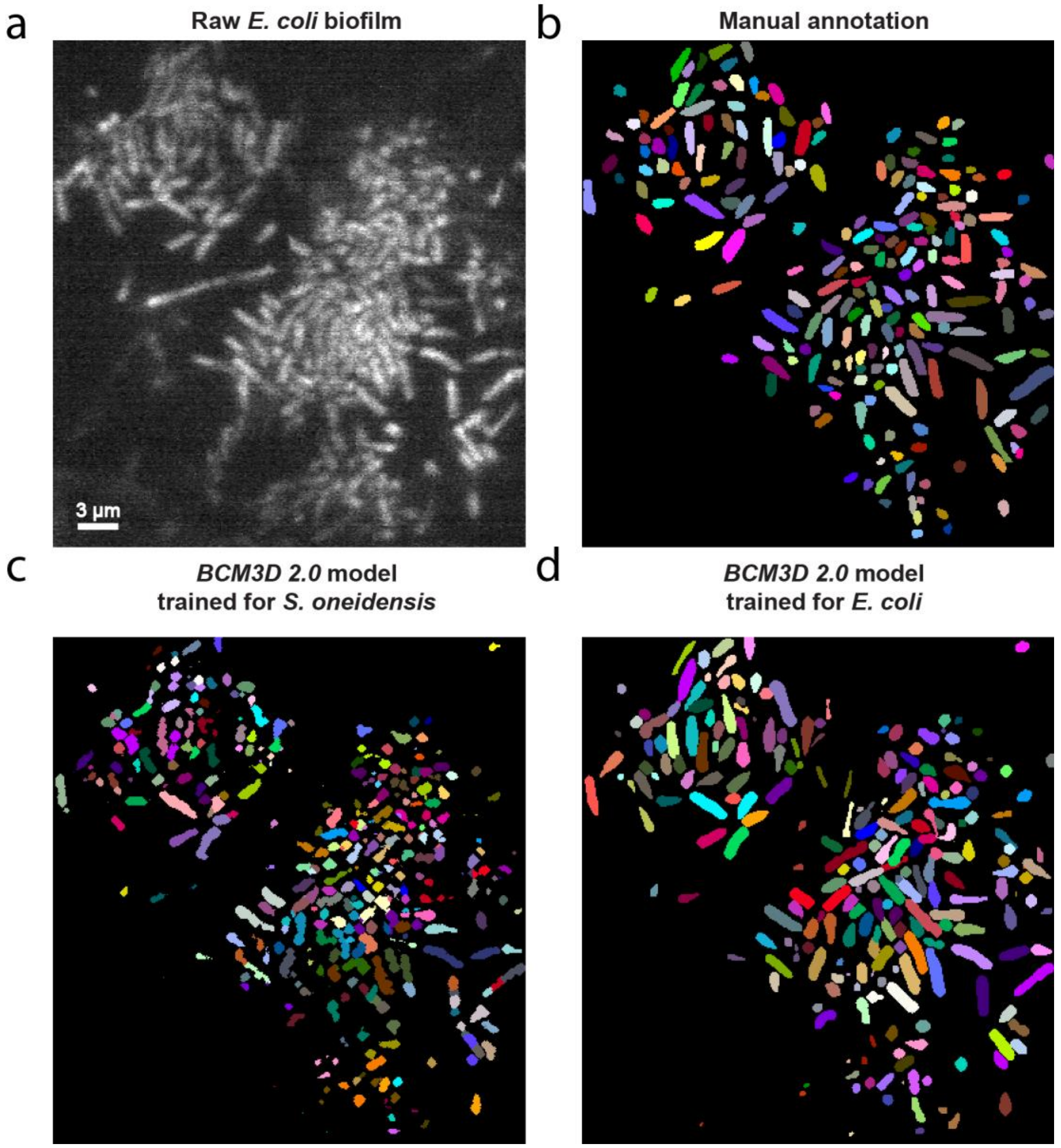


**Figure S1.** **Segmentation results on Zhang et al. 2020 *t* = 600 min data using different training data simulation parameters.**[1] (a) Same 2D cross section shown in Zhang et al. 2020 **Figure 4** t = 600 min. Noted the images have been rotated 90 degrees and flipped horizontally in this figure. (b) Manual annotation result. (c) Segmentation result produced by *BCM3D 2.0* using cell diameter and cell length (d, l) parameters (0.6 µm, 2 µm) in simulation. (d) Segmentation result produced by *BCM3D 2.0* using cell diameter and cell length (d, l) parameters (1 µm, 3µm)

in simulation.

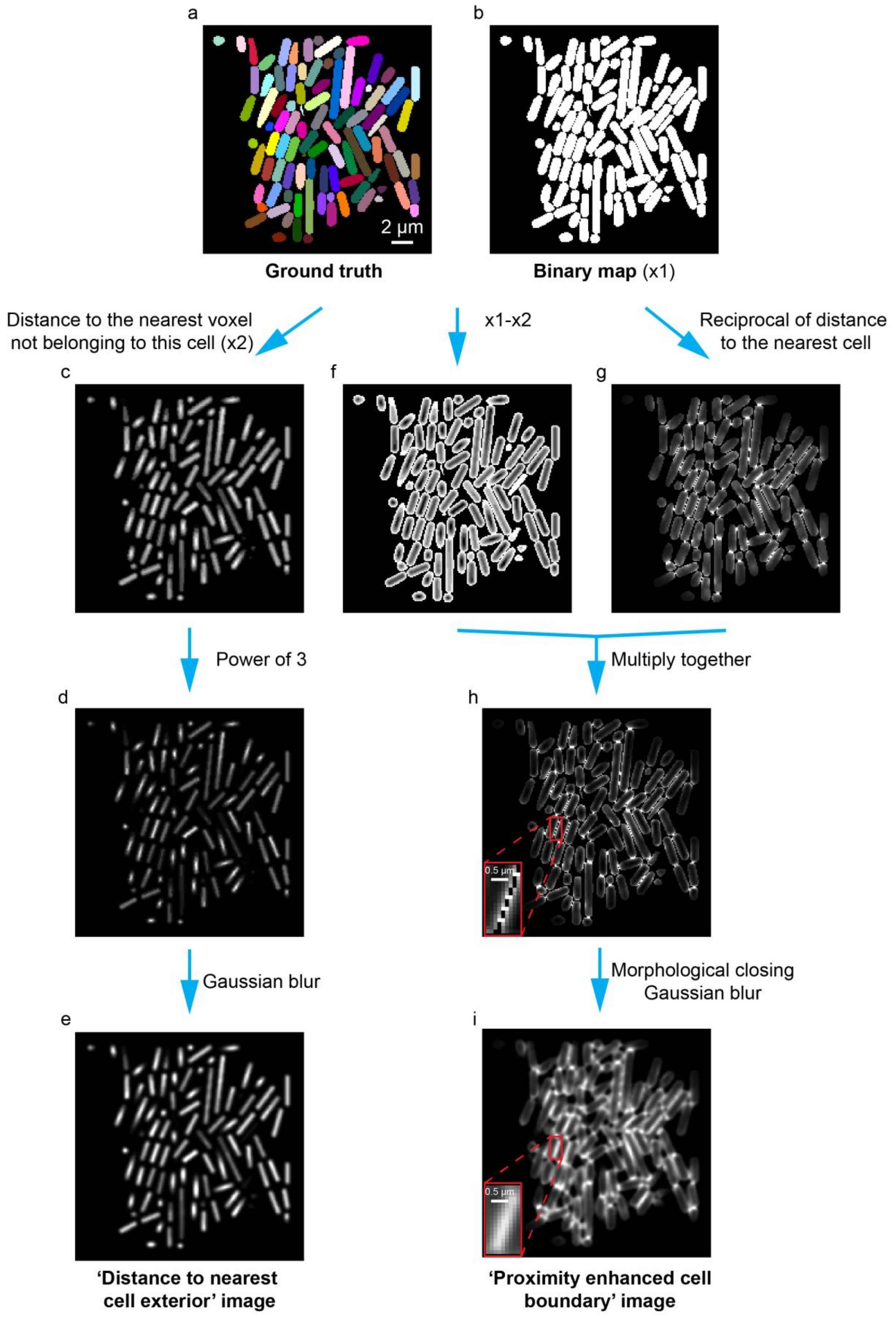
a
b
2 µm
Ground truth
Binary map (x1)
Distance to the nearest voxel
not belonging to this cell (x2)
x1-x2
Reciprocal of distance
to the nearest cell
c
f
g
Power of 3
Multiply together
d
h
0.5 µm
Gaussian blur
Morphological closing
Gaussian blur
e
i
0.5 µm
'Distance to nearest
cell exterior' image
'Proximity enhanced cell
boundary' image

**Figure S2. Schematic of generating intermediate image representations.** (a) Ground truth cell positions. (b) Binary maps based on the ground truth. (c) Images of distance to the nearest voxel not belonging to this cell. (d) A steeper gradient in distance values is obtained by raising each voxel value in panel c to the third power. (e) Smooth images to get 'distance to nearest cell exterior' image representation. (f) Obtain cell boundary by subtracting (c) from (b). (g) Highlight boundary areas that are close to other cells by calculating reciprocal of distance to the nearest cell. (h) Multiply (f) and (g). The inset shows small holes between two cells' boundary. (i) Small holes (inset) are removed in the 'proximity enhanced cell boundary' image by morphological closing and Gaussian blurring.

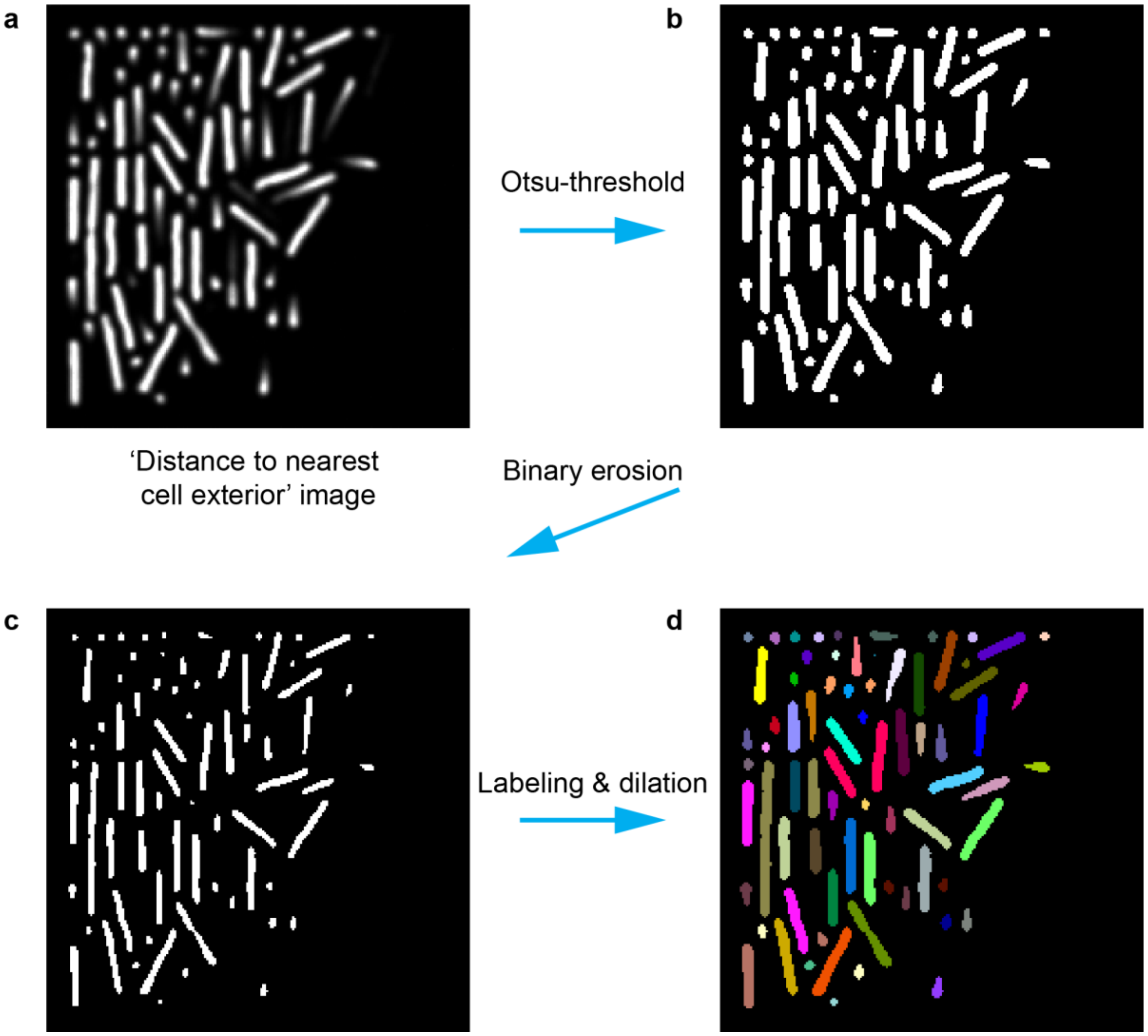


**Figure S3. Post-processing step 1: Thresholding of CNN-produced 'distance to nearest cell exterior' image.** (a) CNN-produced 'distance to the nearest cell exterior' image. (b) Apply Otsu-threshold to obtain binary images. (c) To split clusters that are only connected by one or two voxels, the boundary voxels of each object were set to zero (binary erosion). (d) Identify individual cell objects by labeling connected regions and then add back boundary voxels that were erased in panel c.

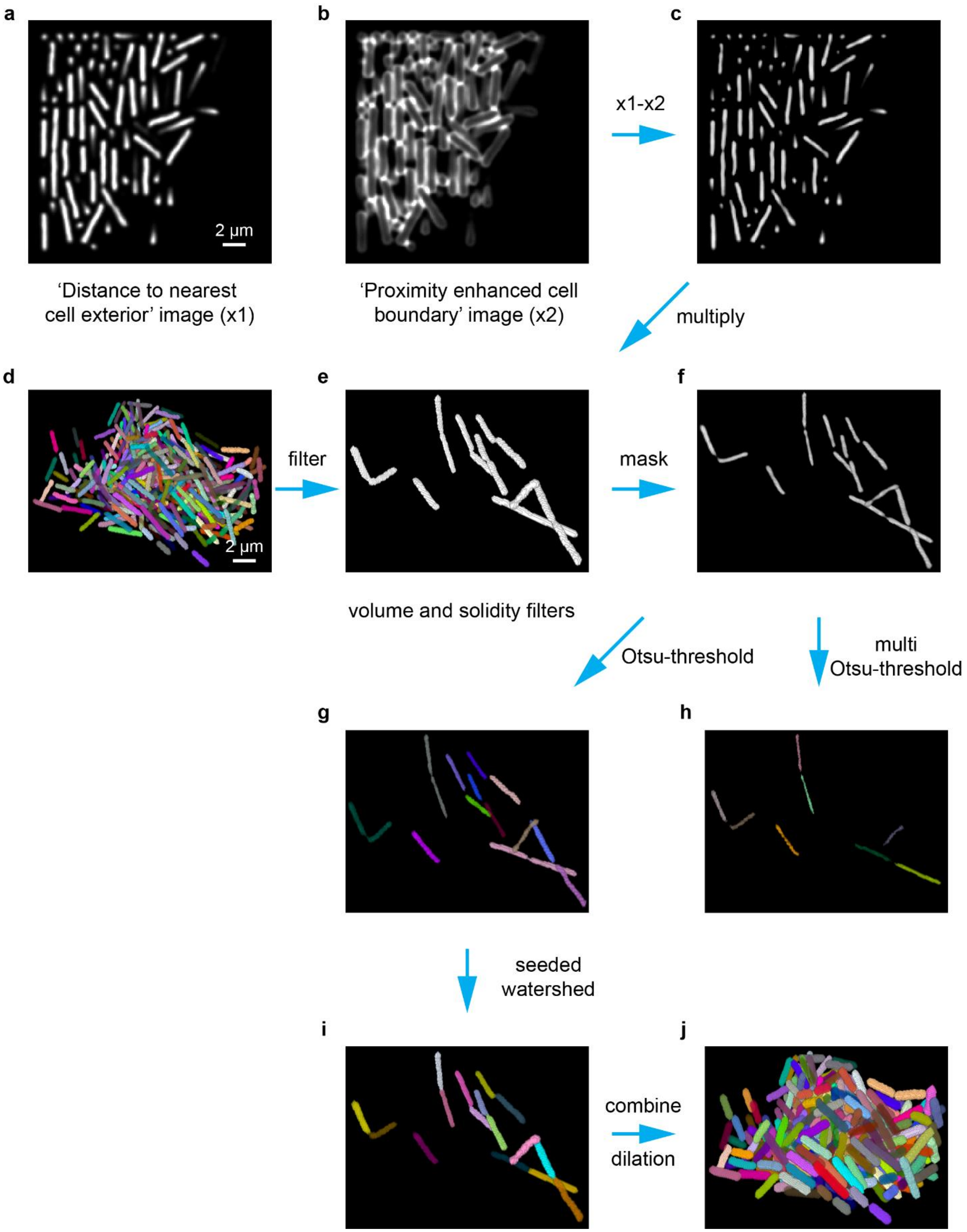
a
b
c
2 µm
x1-x2
‘Distance to nearest cell exterior’ image (x1)
‘Proximity enhanced cell boundary’ image (x2)
multiply
d
e
f
2 µm
filter
mask
volume and solidity filters
Otsu-threshold
multi Otsu-threshold
g
h
seeded watershed
i
j
combine
dilation

48

49

**Figure S4. Post-processing step 2: Combining of intermediate image representations.** (a) CNN-produced ‘distance to nearest cell exterior’ image. (b) CNN-produced ‘proximity enhanced cell boundary’ image. (c) Generate difference map by subtracting (b) from (a) and set negative values to zero. (d) Segmentation results from the process describe in **Figure S3**. (e) Identify objects that need further processing using volume and solidity filters (see Methods). Identified objects are outlined in a binary mask. (f) Mask image in panel c by multiplying it with image in panel e. (g) To split under-segmented clusters, apply seeded-watershed to image in panel f. Seeds are obtained by applying Otsu-thresholding to the image in panel f. (h) If there still are under-segmented clusters in the image in panel g, additional seeds are obtained by applying multi-level Otsu-thresholding the image in panel f. (i) Combine segmented objects from images in panels g and h. (j) Combine (i) and (d) to get final segmentation results.

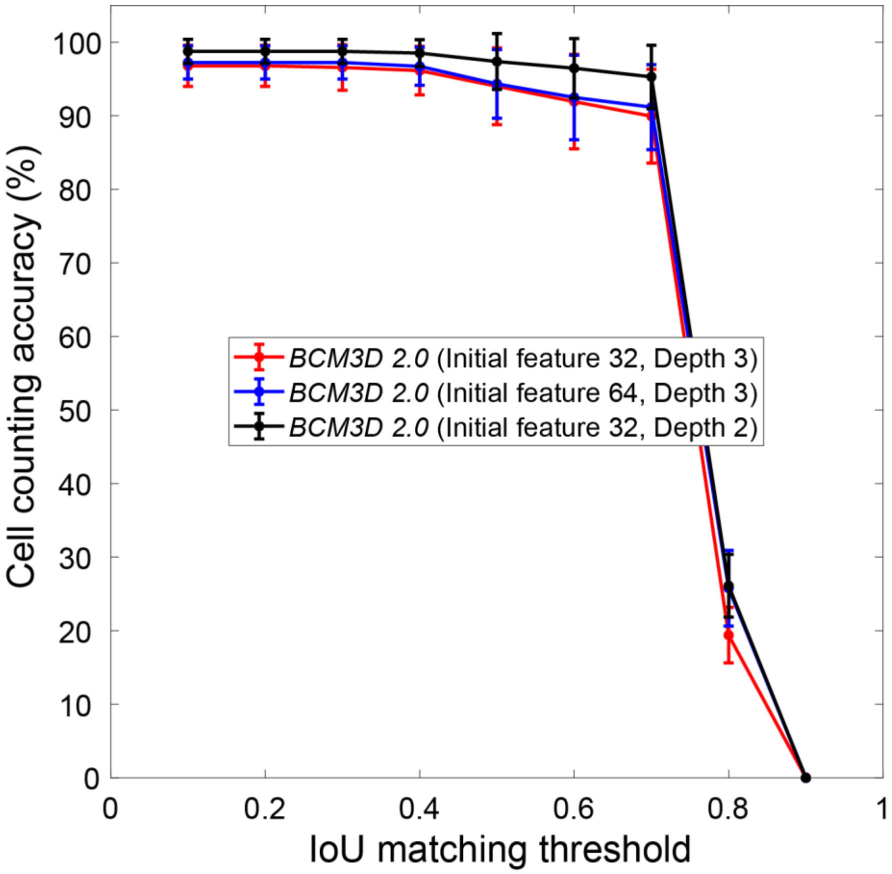


**Figure S5. Comparison of segmentation accuracies achieved by BCM3D 2.0 using different hyperparameter combinations.** The networks were trained with U-Net backbone depth 3, 32 initial features, U-Net depth 3, 64 initial features, and U-Net depth 2, 32 initial features respectively. Segmentation accuracy is parameterized in terms of cell counting accuracy (y axis) and IoU matching threshold (x axis). Each data point is the average of N=10 independent biofilm images. Data are presented as mean values ± one standard deviation.

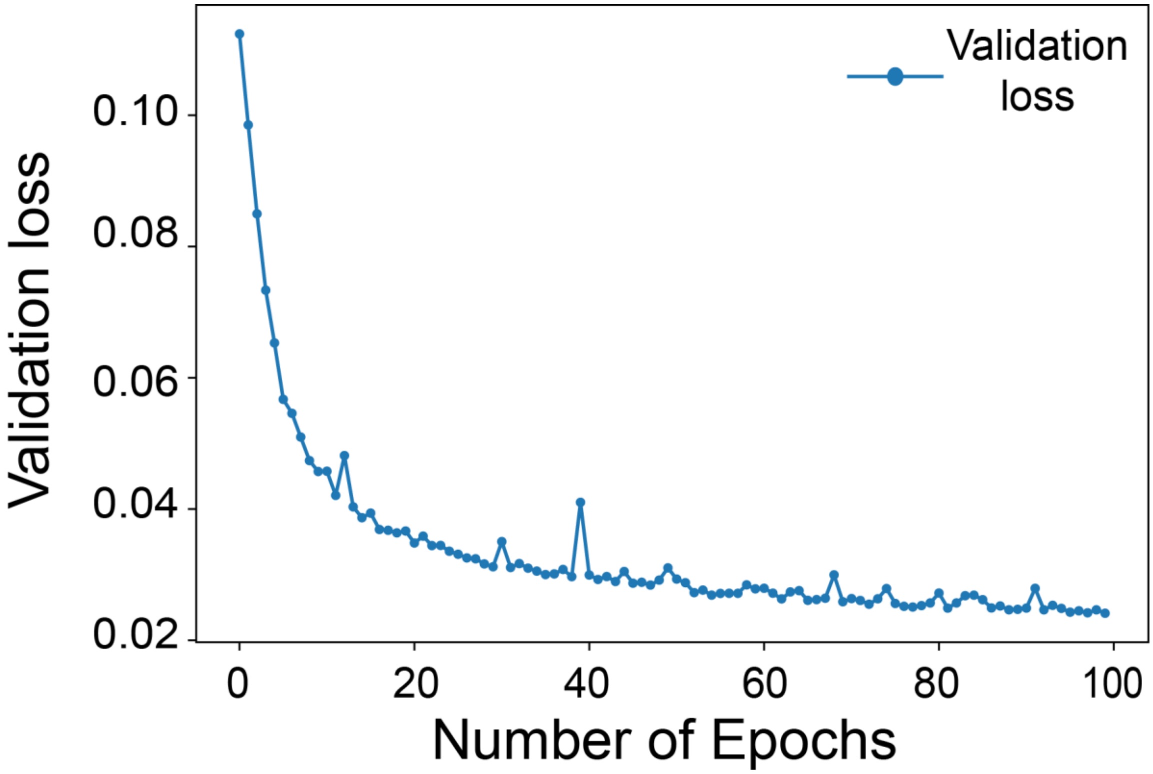


**Figure S6. Validation loss over number of epochs in training**. A U-Net network architecture depth of 2, a convolution kernel size of 3, 32 initial feature maps was trained for 100 epochs. Twenty simulated biofilm images with randomly selected cell densities and signal-to-background ratio were used to train the model.

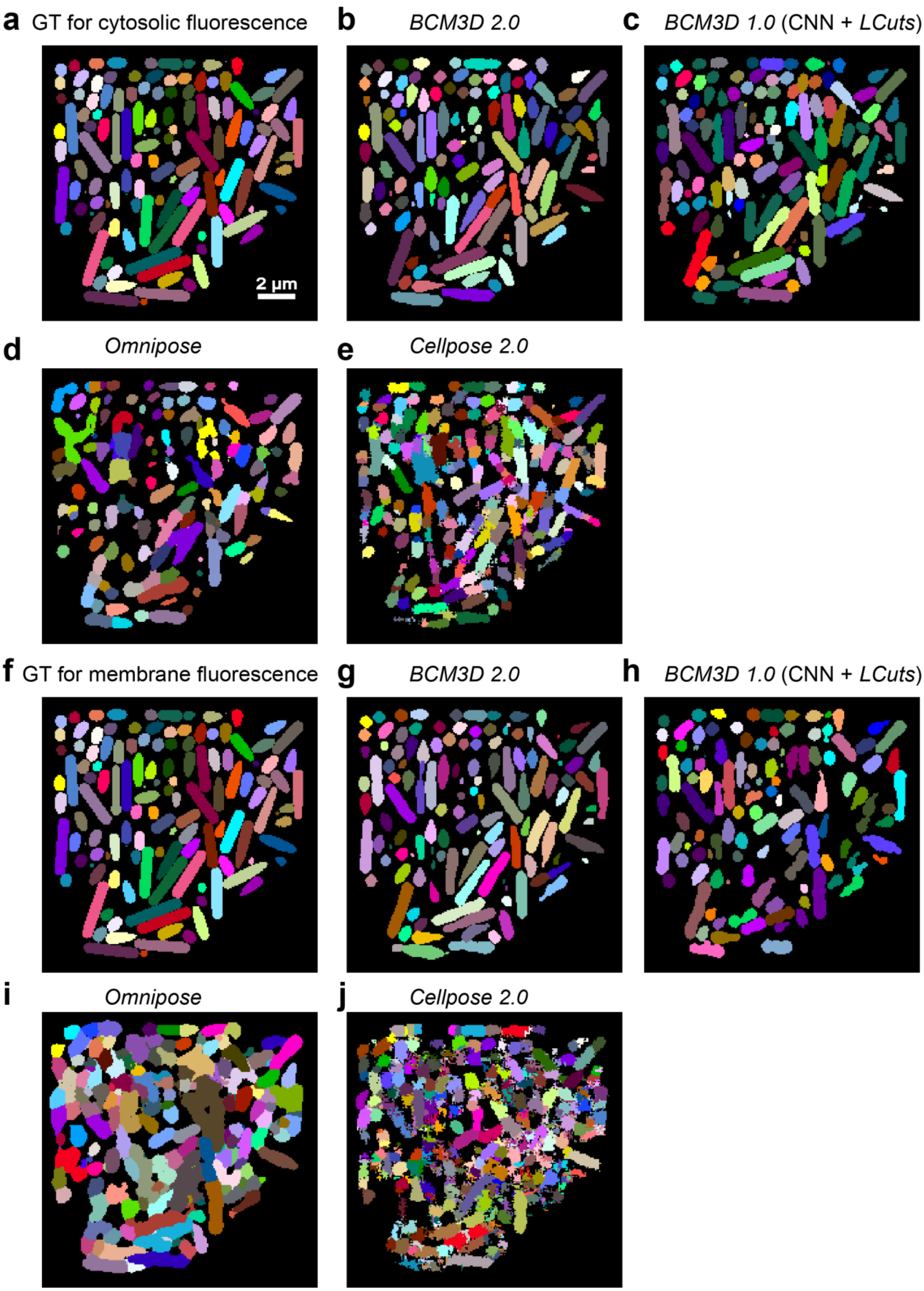
a GT for cytosolic fluorescence
b BCM3D 2.0
c BCM3D 1.0 (CNN + LCuts)
2 µm
d Omnipose
e Cellpose 2.0
f GT for membrane fluorescence
g BCM3D 2.0
h BCM3D 1.0 (CNN + LCuts)
i Omnipose
j Cellpose 2.0

**Figure S7 Visualization of segmentation results for simulated datasets used for Figure 2e and 2f.** (a, f) 2D cross section through the ground truth cell volumes used to generate the simulated images in **Figure 2b** and **2d**. (b-e) Segmentation results produced by *BCM3D 2.0*, *BCM3D 1.0* (CNN+*LCuts*), *Omnipose*, and *Cellpose 2.0* respectively for cytosol-labeled biofilms. (g-j) Segmentation results produced by *BCM3D 2.0*, *BCM3D 1.0* (CNN+*LCuts*), *Omnipose*, and *Cellpose 2.0* respectively for membrane-labeled biofilms.

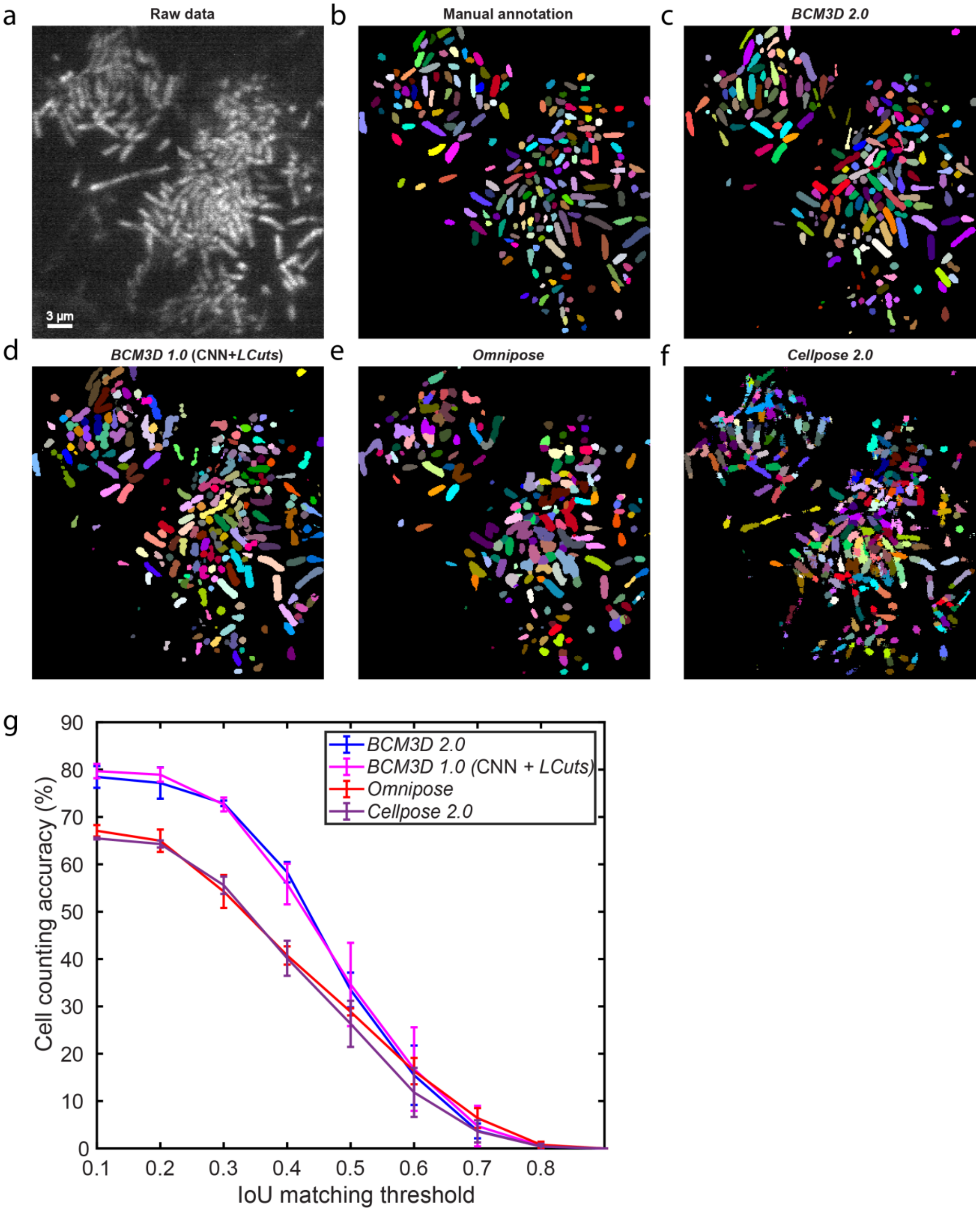


**Figure S8 Comparison of segmentation results on Zhang et al. 2020 *t* = 600 min data.**[1] Noted the images have been rotated 90 degrees and flipped horizontally in this figure. (a) Same 2D cross section shown in Zhang et al. 2020 **Figure 4** t = 600 min. (b) Manual annotation result. (c)

Segmentation result produced by *BCM3D 2.0*. (d) Segmentation result produced by *BCM3D 1.0 CNN + LCuts*. (e) Segmentation result produced by *Omnipose* trained from scratch for up to 800 epochs using the same simulated training dataset that *BCM3D 2.0* was trained on. (f) Segmentation result produced by *Cellpose 2.0* fine-tuned for up to 250 epochs using the randomly selected 2D slices of the same simulated training dataset that *BCM3D 2.0* was trained on. (g) Segmentation accuracy is parameterized in terms of cell counting accuracy ($y$-axis) and IoU matching threshold ($x$-axis). Each data point is the average of the cell counting accuracies calculated using annotation maps traced by $N = 3$ researchers. Data are presented as mean values ± one standard deviation indicated by error bars.

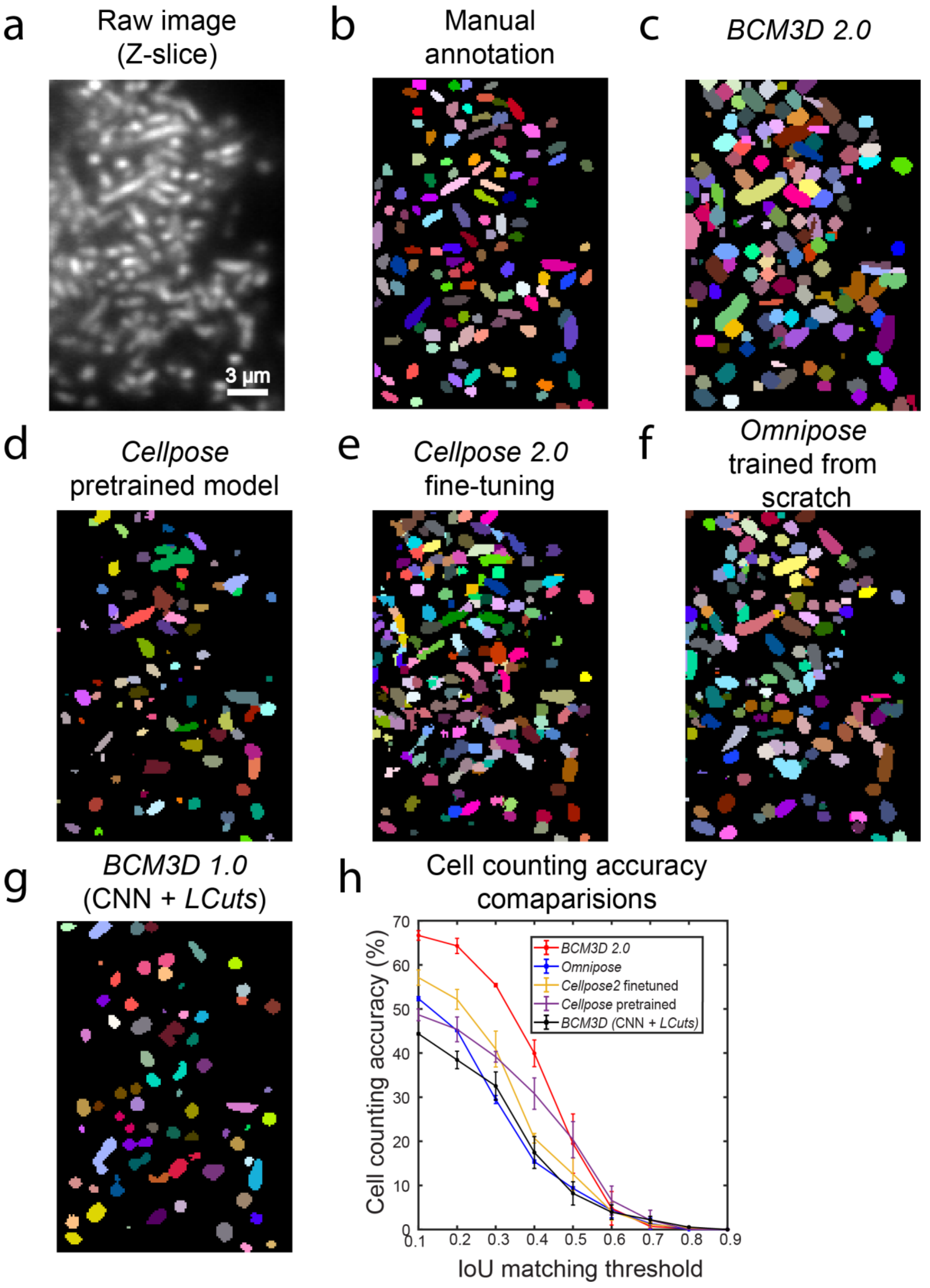
a
Raw image
(Z-slice)
3 μm
b
Manual
annotation
c
BCM3D 2.0
d
Cellpose
pretrained model
e
Cellpose 2.0
fine-tuning
f
Omnipose
trained from
scratch
g
BCM3D 1.0
(CNN + LCuts)
h
Cell counting accuracy
comaparisions
Cell counting accuracy (%)
IoU matching threshold
BCM3D 2.0
Omnipose
Cellpose2 finetuned
Cellpose pretrained
BCM3D (CNN + LCuts)
0
10
20
30
40
50
60
70
0.1
0.2
0.3
0.4
0.5
0.6
0.7
0.8
0.9

**Figure S9 Performance comparisons on *S. oneidensis* dataset in Figure 3.**

(a) Small 2D cross section selected from the large 3D biofilm shown in **Figure 3**. (b) Manual annotation result. (c) Segmentation result produced by *BCM3D 2.0*. (d) Segmentation result produced by the pretrained *Cellpose* model. (e) Segmentation result produced by *Cellpose2* fine-tuned using the simulated training data from this work. Cellpose2 can be used for quickly prototyping new specialist models from a generalist model with minimal new training data. We note that *Cellpose2* only uses 2D data for training and fine-tuning, so thirty different 2D slices in all three dimensions of the 3D training data were selected to fine-tune the network. (f) Segmentation result produced by *Omnipose* trained from scratch for up to 800 epochs using the same simulated training dataset that *BCM3D 2.0* was trained on. Validation loss was observed to stop decreasing after approximately 600 epochs. (g) Segmentation result produced by *BCM3D 1.0 CNN + LCuts* (h) Segmentation accuracy is parameterized in terms of cell counting accuracy ($y$-axis) and IoU matching threshold ($x$-axis). Each data point is the average of the cell counting accuracies calculated using annotation maps traced by $N = 2$ researchers. Data are presented as mean values $\pm$ one standard deviation indicated by error bars.

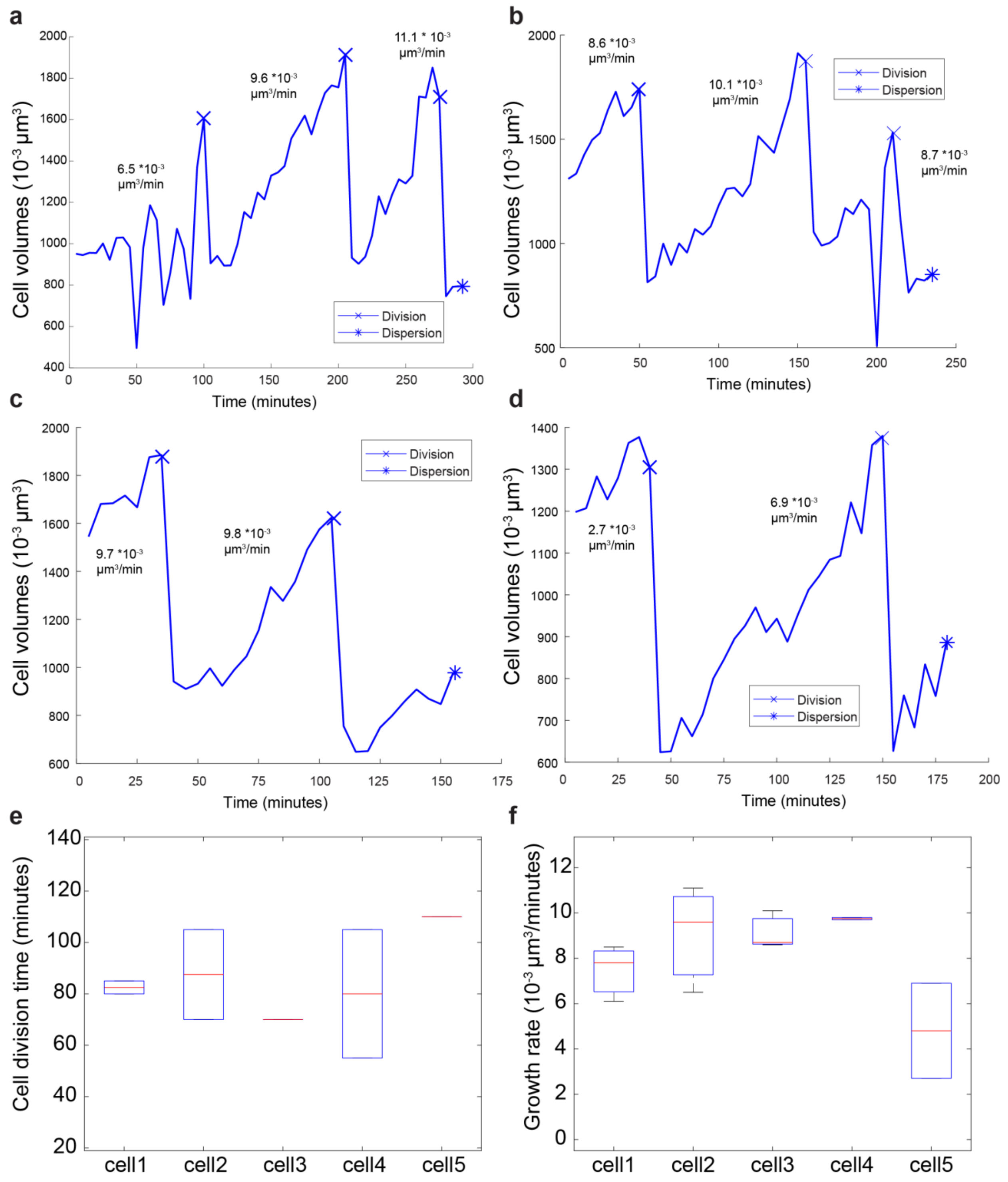


**Figure S10.** (a-f) Cell volumes over time for four additional cells. The solid line represents a manually annotated trajectory where cell division and dispersion events are detected manually and indicated on the trajectories. Single-cell growth rate was measured by calculating the slope of a line that connects both the beginning and end points for each trajectory. (e) Box plot of cell division time for the selected fives cell (four in this figure plus one in Figure 5). (f) Box plot of single-cell growth rate for the selected five cells.

**Thresholding of CNN-produced 'distance to nearest cell exterior' images (Figure S3)**

Thresholding of CNN-produced 'distance to nearest cell exterior' images is the first step of the post-processing pipeline. Predicted 'distance to nearest cell exterior' images were first normalized by a simple percentile-based normalization method (**Figure S3a**), which we define for an input u as

$$N(u; p_{low}, p_{high}) = \frac{u - perc(u, p_{low})}{perc(u, p_{high}) - perc(u, p_{low})}$$

Where perc(u,p) is the p-th percentile of all voxel values of u. We typically use $p_{low} = 3$ and $p_{high} \in (99.5, 99.9)$. After applying Otsu-thresholding to the 'distance to nearest cell exterior' image to obtain a binary image (**Figure S3b**), connected voxel clusters can be isolated and identified as single cell objects by labeling connected regions. To split clusters that are only connected by one or two voxels, the boundary voxels of each object were set to zero before labeling connected regions (**Figure S3c**). After labeling, the erased boundary voxels were added back to each object (**Figure S3d**). A conservative size-exclusion filter was applied: small objects with volume smaller than the radius cubed of the targeted cells were considered background noise and filtered out.

**Post-processing of initial thresholding result (Figure S3) by introducing 'proximity enhanced cell boundary' images (Figure S4)**

Thresholding of the 'distance to nearest cell exterior' image produces a binary image (background = 0, cell = 1), where groups of connected, non-zero voxels identify individual cells in most cases. However, when cells are touching, they are often not segmented as individuals, but remain part of the same voxel cluster (undersegmentation). On the other hand, a small part of a cell may be erroneously identified as another cell (oversegmentation). To address these errors and further improve the segmentation accuracy, we included ‘proximity enhanced cell boundary’ image, seeded watershed, and multi-level Otsu thresholding to post-process the binary images obtained from the normalized ‘distance to nearest cell exterior’ images (**Figure S4**).

Step 1. Objects that need further processing were found by evaluating its volume and solidity, i.e., the volume to convex volume ratio. Here, volume is defined as the number of voxels occupied by an object. Convex volume is defined as the number of voxels of a convex hull, which is the smallest convex polygon that encloses an object. The upper limit was found by using the interquartile rule, i.e. the upper limit is quartile 3 (Q3) plus 1.5 times interquartile range (IQR). If an object's volume or solidity is larger than the upper limit, it will be singled out for further processing. All these objects together generate a new binary image (**Figure S4e**).

Step 2. To identify and delineate individual cells in the undersegmented connected voxel clusters, CNN-produced ‘proximity enhanced cell boundary’ images were first normalized by the same percentile-based normalization method. We used seeded watershed after combining ‘distance to nearest cell exterior’ images and ‘proximity enhanced cell boundary’ images. Specifically, we generated a difference map by subtracting the ‘proximity enhanced cell boundary’ image from the ‘distance to nearest cell exterior’ image and then set all negative valued voxels to zero (**Figure S4a, b, c**). This difference map was then multiplied by the binary image generated in Step 1 (**Figure S4f**). Then, the resulting image was segmented by seeded watershed. Seeds were obtained by Otsu thresholding of the difference map and seeds with a volume smaller than 30 voxels were removed (**Figure S4g**). These new objects were again evaluated by their volume and solidity; if there still exist unmatched objects, a multi-level Otsu thresholding will be applied to further generate seeds. Unlike simple Otsu thresholding, which only separates voxels into two classes, foreground and background, multi-level Otsu calculates several thresholds, determined by the number of desired classes. Here, we used the following five classes: background, transition area between background and cell border, cell border, transition area between cell border and cell interior, cell interior. Seeds were extracted by using the third and the fourth threshold successively, i.e. the threshold between the third and the fourth classes (cell border, transition area between cell border and cell interior), and the threshold between the fourth and the fifth classes (transition area between cell border and cell interior, cell interior). The same size filter was used to remove unreasonable small seeds (**Figure S4h**).

Step 3: All post-process objects were added together (**Figure S4i**) and combined with the initially reasonable objects, i.e., the objects haven't been singled out in Step 1. A conservative size-exclusion filter was applied: small objects with volume 10 times smaller than the upper limit volume were considered unreasonable small parts and filtered out. Since the 'distance to nearest cell exterior' images were confined to the cell interior, we dilated each object by 1-2 voxels to increase the cell volumes using standard morphological dilation (**Figure S4j**).
